# Active control of surface immobilization in the foraging behavior of *Paramecium*

**DOI:** 10.64898/2026.09.19.752847

**Authors:** Ali Hosseini, Wilder Boyden, Célia Fosse, Marcel Stimberg, Robert D. Guy, Romain Brette

**Affiliations:** Sorbonne Université, CNRS, Institute of Intelligent Systems and Robotics (ISIR), 75005 Paris, France; Department of Mathematics, University of California, Davis, CA 95616

## Abstract

*Paramecium* is a common freshwater ciliate, a unicellular swimming eukaryote that feeds on bacteria. Since the discovery that its motility is controlled by action potentials, which earned it the nickname of “swimming neuron”, many studies have documented the effect of various stimuli on its behavior, as well as their physiological basis, highlighting the richness of adaptive behavior in this unicellular organism. The logic of its autonomous behavior has received comparably less attention.

For example, when *Paramecium* encounters a surface, it may slide or avoid it by a directional change, triggered by an action potential, or it may immobilize on the surface, a behavior often called *thigmotaxis*. Here we ask why and how *Paramecium* immobilizes on surfaces. With long term behavioral tracking, we show that this variable behavior is a component of *Paramecium*’s foraging behavior in the presence of bacteria, where the organism alternates between exploring and stopping on surfaces to feed. This immobilization is physiologically controlled by a calcium influx. Using mutant strains, high speed imaging, particle image velocimetry and hydrodynamics simulation, we further show that surface immobilization occurs by an inhibition of locomotor cilia, while oral cilia continue beating strongly.

These findings open the perspective of studying the physiological control of foraging in a unicellular organism, a complex ecologically relevant behavior potentially involving sensory processing, memory, decision making, and motor control.

## Introduction

*Paramecium* is a common freshwater ciliate, a unicellular eukaryote that swims by beating its thousands of cilia and feeds on bacteria by filtering the water. Because of its abundance and large size (up to 300 µm), its rich behavior was already documented in detail in the 19^th^ century^1^. In the 1960s, it was found that its motility is electrically controlled by action potentials, and *Paramecium* effectively became a model organism in neuroscience under the nickname of “swimming neuron”^2^.

Most studies of *Paramecium* behavior have focused on the influence of various stimuli on its swimming behavior. For example, mechanical stimulation of its anterior end triggers an “avoiding reaction” mediated by a calcium-based action potential, where the organism swiftly swims backward then changes direction^3^. However, this reaction is not systematic: in its spontaneous swimming behavior, *Paramecium* often slides on surfaces without changing direction^4^, or reacts with a delay^5^. In fact, in its natural environment or in culture vessels, *Paramecium* is often found immobile on surfaces. As Jennings noted already in 1897, “*the large majority of [paramecia] will generally be found clinging to some solid body. A large number are attached to the decaying plant tissue just below the surface of the water*”^6^, a phenomenon he termed *thigmotaxis*.

This is intriguing because the same stimulus appears to induce a variety of behaviors, perhaps depending on context or history. Indeed, a number of studies have shown that ciliate behavior can change with previous interactions^7^, through habituation^8–10^ or conditioning^2,11,12^, or with environmental context^13–15^. However, why and how *Paramecium* sometimes immobilizes on surfaces is unknown. Kitamura linked this behavior to sexual behavior (conjugation), where two cells of compatible mating types join and exchange genetic material, and proposed that attachment is mediated by changes in the stickiness of cilia with sexual maturation^16^. However, Iwatsuki *et al.* later found that attachment can also be induced in non-sexually reactive cells, simply by increasing the concentration of calcium or potassium in the medium^17,18^. Previously, Saunders proposed that *Paramecium* attaches by projecting trichocysts^19^, which are needle-like organelles anchored below the membrane, released by calcium-controlled exocytosis^20^. Immobilization was also mentioned as a transient “inactive” phase during equilibration in an inorganic solution^21^.

Here we ask why and how *Paramecium* immobilizes on surfaces, using long term behavioral tracking, pharmacology, mutant strains, high speed imaging, particle image velocimetry and hydrodynamics simulation. Starting from the observation of spontaneous behavior in the culture medium, we first show that in the presence of bacteria, *Paramecium* alternates between fast swimming and immobilization on surfaces where it feeds, much like the foraging behavior of animals. In this context, behavioral variability is functional, as it allows exploration. We then show that immobilization is physiologically mediated by a calcium influx. *Paramecium* then immobilizes on surfaces not by sticking or anchoring, but by inhibiting its locomotor cilia, while oral cilia continue beating strongly.

## Results

### Foraging behavior of *Paramecium*

In all experiments, we used the homozygous strain 51 of *P. tetraurelia,* with a culture protocol ensuring that cells have similar clonal age and are not autogamous (see Methods). In the culture tube, immobile paramecia are routinely seen on the surfaces of the tube as well as on top, where a biofilm often forms (Fig. 1A). Careful observation reveals that this picture is dynamic, as cells can repeatedly attach and detach from the biofilm (Video S1). To study this phenomenon in more detail, we place a drop of bacterized culture medium with growing paramecia (logarithmic phase) between two glass slides spaced by 520 µm. After a few minutes, many cells appear to be immobile against the slides, both on top and at the bottom (Fig. 1B). Since *Paramecium* is about 4% heavier than water^22^, this indicates that cells did not simply stop swimming and fall. Indeed, on closer inspection, we observe intense ciliary activity and a flux of bacteria moving in the anteroposterior direction, especially near the oral groove (Video S2). Inside the cells, we distinguish many slowly moving food vacuoles, indicating active digestion. These observations show that immobile cells are not inactive, but actively feeding.

**Figure 1.**
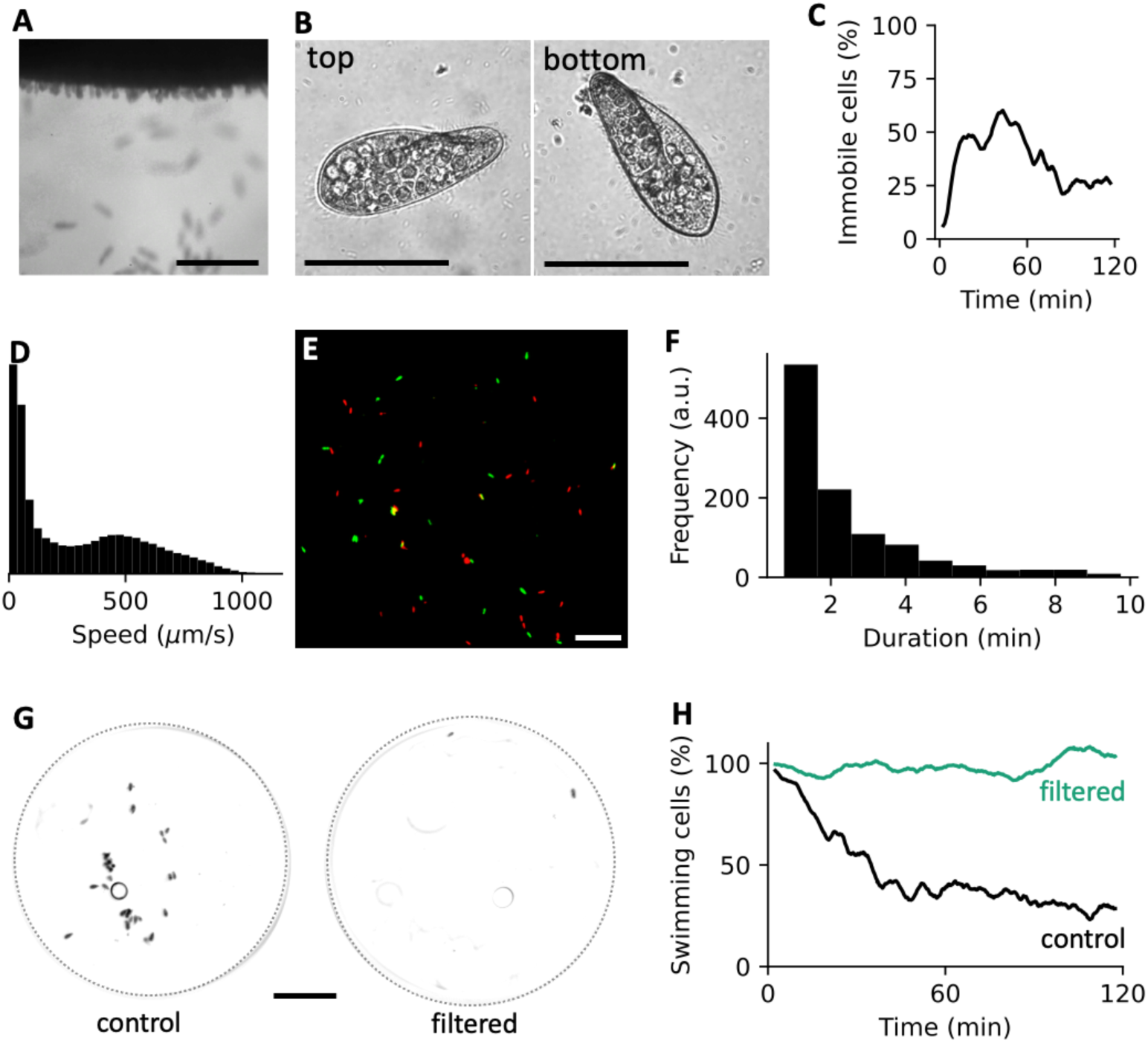
Thigmotaxis in the culture medium. A, Paramecia attached to the surface biofilm and walls of the culture tube (0.5x; scale bar: 1 mm). B, Examples of cells attached to the top and bottom glass slides in a drop of culture medium (40x, scale bar: 100 µm). C, Proportion of immobile cells vs. time in the culture medium drop (n = 55). D, Speed distribution of swimming cells at t = 1 h (measured over 5 min). E, Immobile cells at t = 60 min (red) and t = 65 min (green; overlaps appear yellow). F, Distribution of attachment duration. G, Immobile cells at t = 1 h in the control bacterized drop and in the filtered drop (dotted circles: outlines of the drops). H, Proportion of swimming cells in the control bacterized drop (black, n = 24) and in the filtered drop (green, n = 38), relative to the initial number of swimming cells.

We then examine the dynamics of attachment. To this end, we observe a drop of growing paramecia (n = 55) at lower magnification (0.5x) with a wide field camera, so as to capture all cells in the entire drop, and we track trajectories for two hours (see Methods). To track swimming cells, we first remove the background, calculated by averaging the frames over a 15 s period (Video S3). This reduces the data bandwidth by a factor of about 100. Cells that remain immobile for more than 15 s appear in the background frames, which are also saved (Video S4). We then count immobile and swimming cells.

Within 20 min, the proportion of immobile cells rises over 50% (Fig. 1C), then progressively settles to about 25%. When we measure the speed of swimming cells in a 5 min interval around *t* = 1 h, we observe that there are two modes, one near 0 µm/s and another one near 500 µm/s (Fig. 1D). Thus, at any given time, the population is split into fast swimming cells and very slow or immobile cells. Furthermore, the positions of immobile cells change completely over a 5 min interval (Fig. 1E). When we track cells in background frames (i.e., immobile for more than 15 s), we find that they stay immobile for a variable duration of around 3 min (193 ± 271 s, mean ± s.d.), distributed approximately exponentially (Fig. 1F). Thus, cells appear to alternate between fast swimming and transient immobilization on surfaces where they feed, similar to the foraging behavior of many animals.

We then compare swimming behavior in the presence and absence of bacteria. To this end, we pick two groups of cells from the same culture tube in early logarithmic phase (high bacterial density) and place one group in the original medium, and the other one in the same medium, but with bacteria removed by filtering. We then observe their swimming behavior simultaneously for two hours. The difference is striking (Fig. 1G-H, Video S5): while paramecia progressively stop and even form clusters in the bacterized medium, almost all paramecia swim in the filtered medium (here we report the number of swimming cells because cell clustering makes the immobile cell count unreliable).

Next, we show that immobilization is under physiological control.

### Surface immobilization requires a calcium influx

Calcium signaling has been identified in several important behaviors of *Paramecium*: the avoiding reaction depends on L-type calcium voltage-gated channels^3,23^, the escape reaction depends on a hyperpolarization-activated calcium current^24^, and trichocyst release involves extracellular calcium and calcium-induced calcium release^20^. We then tested whether extracellular calcium is necessary for surface immobilization. A difficulty is that *Paramecium* does not survive in the absence of calcium, and the concentration of calcium in the culture medium is not precisely known. Therefore, we observed the effect of different concentrations of the calcium chelator EGTA applied to the bacterized culture medium (Fig. 2A). With 0.32 mM EGTA, cells immobilized similarly as in the control (green vs. black), reaching nearly 50% of immobilized cells after an hour. With 0.37 mM EGTA, only 18% of cells immobilized (Fig. 2A, orange; Fig. 2B). With 0.4 mM EGTA, all cells died. Thus, extracellular calcium is required for immobilization, above the concentration necessary for surviving.

**Figure 2.**
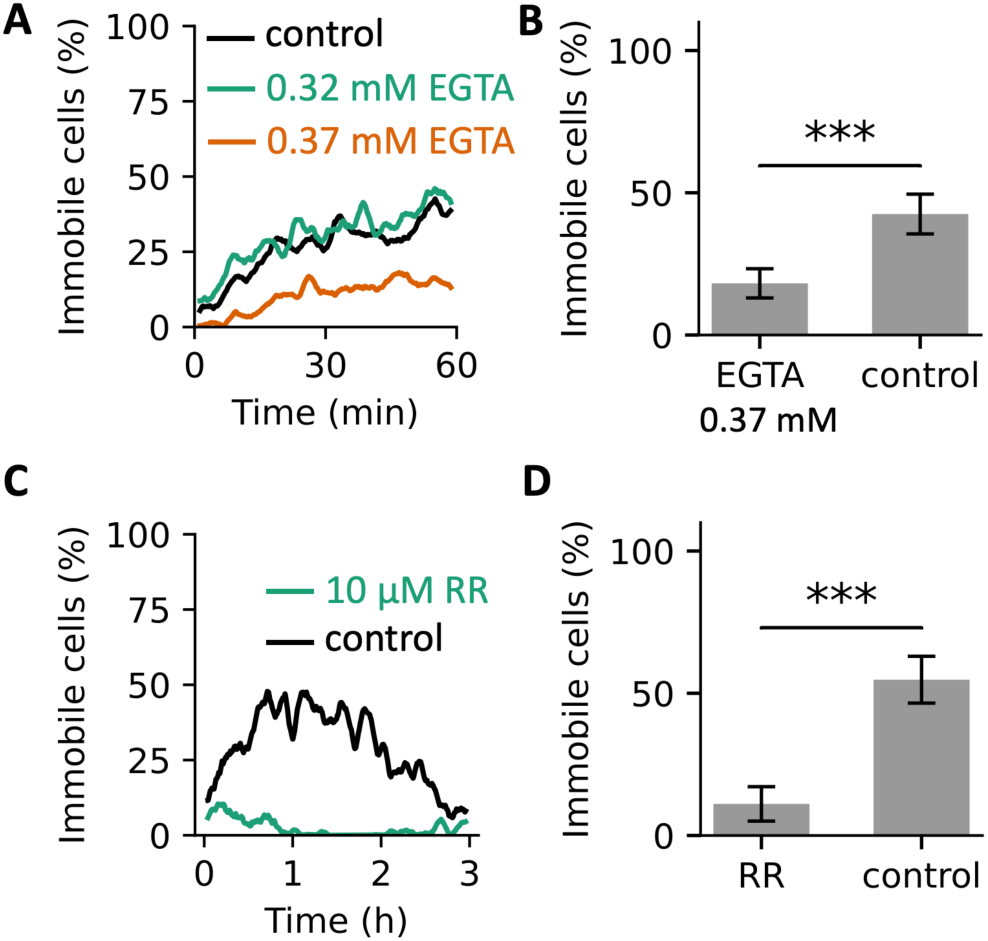
Calcium dependence of immobilization. A, Proportion of immobile cells vs. time in the bacterized culture medium (black, n = 56), and with 0.32 mM EGTA (green, n = 44) and 0.37 mM EGTA (orange, n = 51). B, Maximum proportion of immobile cells in the 0.37 mM EGTA and control solution (error bars = s.e.m.; p = 3·10^-8^, Welch’s t-test). C, Proportion of immobile cells vs. time in the bacterized culture medium with added 10 µM ruthenium red (green, n = 30) vs. control (black, n = 38). D, Maximum proportion of immobile cells in the 10 µM RR and control solution (error bars = s.e.m.; p = 6·10^-13^, Welch’s t-test).

We then applied the calcium channel blocker ruthenium red (RR) in the bacterized culture medium. We observed that adding 10 µM of RR almost completely blocked surface immobilization (Fig. 2C, D). In addition, we observed that cell growth was also strongly inhibited by 10 µM RR: cell count increased by about 11% over the 3 h of the experiment vs. 34% in the control.

We conclude that surface immobilization involves a calcium influx, possibly through a TRP channel (one of the main targets of RR). Next, we ask how *Paramecium* immobilizes against surfaces, especially as it can stop on both top and bottom surfaces. Since bacteria are required for this behavior, one hypothesis is that *Paramecium* might attach to bacterial products, such as biofilms. We address this possibility by reexamining previous reports of inactivity in inorganic solutions.

### Surface immobilization in inorganic solutions

Several authors previously observed that *Paramecium* immobilizes on surfaces after being washed in an inorganic calcium-potassium solution, but did not relate it to feeding behavior. Kitamura observed that sexually reactive cells attach on hydrophobic surfaces when washed in an inorganic solution, and attributed this phenomenon to changes in the stickiness of cilia developing with sexual maturation^16^, in relation with their sexual behavior (conjugation). Iwatsuki *et al.* later found that attachment can also occur on glass, and in non-sexually reactive cells, simply by increasing the concentration of calcium or potassium in the medium^17,18^. In a study of gravitaxis, Machemer et al.^21^ briefly mentioned a transient “inactive” phase during equilibration in an inorganic solution.

We checked whether this washing-induced immobilization is similar to what we observed in feeding behavior. We washed late stationary cells (starved) in a solution of 1 mM CaCl_2_ and 4 mM KCl. To make sure cells carry as few bacteria as possible, we transferred them individually to four clean drops in sequence (see Methods), and imaged the final drop covered with a glass coverslip, for 2 h. In this sterile drop, most cells immobilize (Fig. 3A, black; n= 18). As in the bacterized medium, attached cells were found on both bottom and top surfaces (Fig. 3B), with cilia beating (Video S6). Thus, this appears very similar to the feeding behavior observed in the bacterized medium, except there are no bacteria.

**Figure 3.**
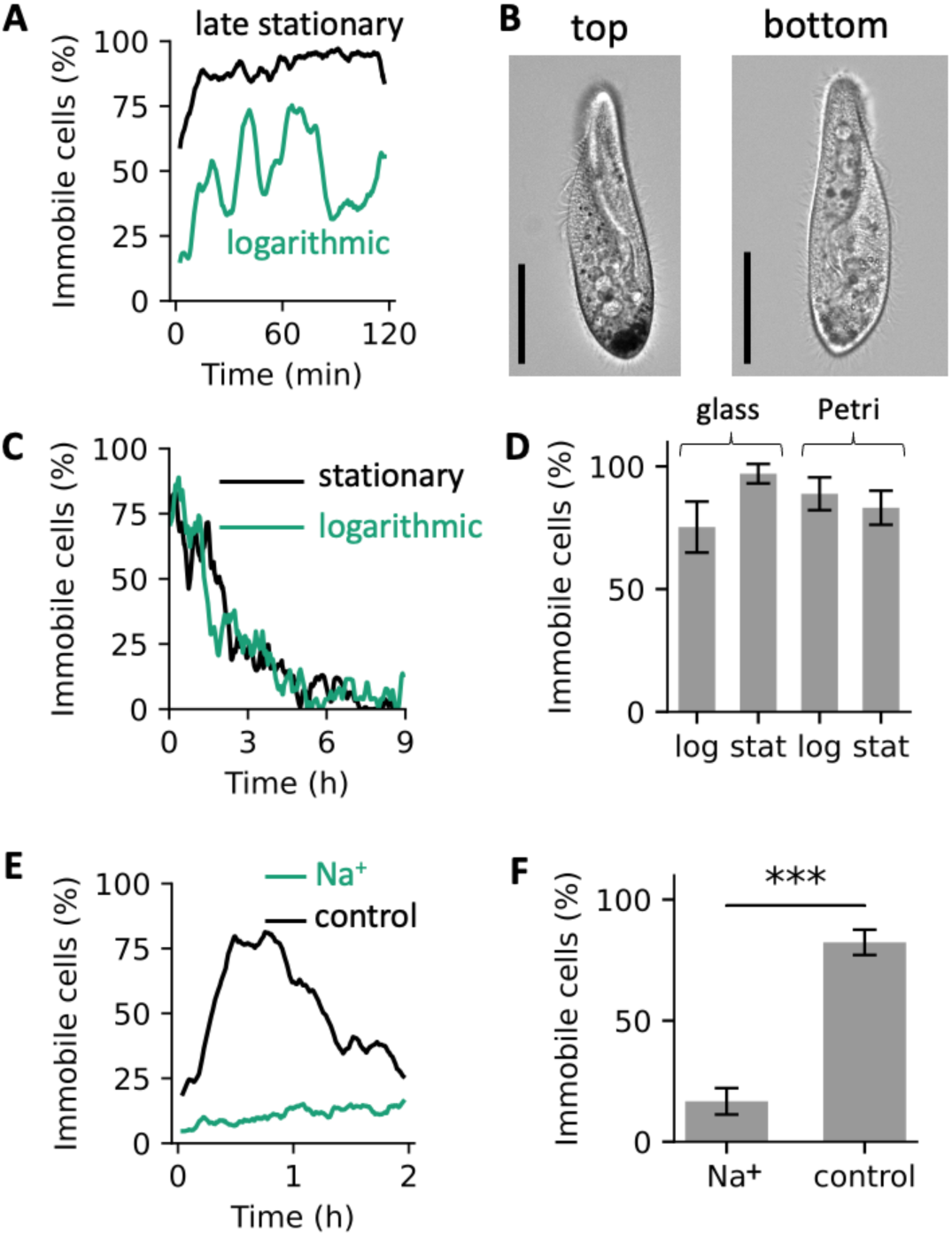
Immobilization in the sterile inorganic medium. A, Proportion of immobile cells vs. time in the inorganic medium drop between glass coverslips, for late stationary cells (black, n = 18) and logarithmic cells (green, n = 19). B, Examples of stationary cells attached to the top and bottom glass coverslips in a drop of inorganic medium (20x, scale bar: 50 µm). C, Proportion of immobile cells vs. time in the inorganic medium drop on a Petri dish, for late stationary cells (black, n = 18) and logarithmic cells (green, n = 19). D, Maximum proportion of immobile logarithmic and stationary cells on glass coverslips and Petri dish (from the data of A and C). Error bars show the standard error of the mean, assuming binomial distributions (i.e., cells are independent). E, Proportion of immobile cells vs. time in the inorganic solution with added 11.5 mM Na^+^ (green, n = 50) vs. control (black, n = 50). F, Maximum proportion of immobile cells in the Na^+^ and control solution (error bars = s.e.m.; p = 5·10^-41^, Welch’s t-test).

We repeated this procedure with growing cells (logarithmic phase) (Fig. 3A, green; n = 19) and many cells also immobilized, although fewer. This is consistent with the observations of Iwatsuki et al.^18^. When glass slides were replaced by a plastic Petri dish, both stationary and logarithmic cells immobilized (Fig. 3C). We then observed the cells for a longer duration (9 h). To our surprise, after a couple of hours, paramecia started to swim again, so that most of them were swimming after 6 h.

Thus, washing paramecia in the sterile inorganic solution induces surface immobilization with the same characteristics as in normal feeding behavior: on both top and bottom surfaces, with strong ciliary beating, and transient. In addition, washing-induced immobilization also requires extracellular calcium and is blocked by ruthenium red^17^. This behavior is seen in both logarithmic and stationary cells and on both hydrophilic (glass) and hydrophobic (Petri dish) surfaces (summarized in Fig. 3D). This shows that paramecia can immobilize on different types of surfaces without attaching to bacterial products.

Yet, we have seen previously that removing bacteria from the culture medium prevents immobilization (Fig. 1H). Why then did cells immobilize in the calcium-potassium solution? We hypothesized that this might be due to the different ionic contents of the two media. The culture medium contains a pH buffer that is rich in Na^+^ (11.5 mM), while there is no Na^+^ in the washing solution. When we added 11.5 mM Na^+^ to the washing solution, immobilization was not induced anymore (Fig. 3E, F). We conclude that surface immobilization is normally triggered by the presence of bacteria, but can also be triggered by depletion of extracellular Na^+^ (which is normally present in fresh water). This means that bacteria act as a signal for immobilization, rather than means of attachment.

*Paramecium* does not immobilize by attaching to bacteria, it can attach and detach spontaneously or pharmacologically, and surface properties are not a critical factor. These observations speak against an important role for ciliary stickiness in this phenomenon. How then does *Paramecium* immobilize against surfaces?

### *Paramecium* immobilizes by inhibiting its locomotor cilia

In 1925, Saunders proposed that *Paramecium* attaches with trichocysts^19^, which are needle-like organelles docked below the membrane, projected by a calcium-based exocytosis apparatus^20^. To test this hypothesis, we examined the *trichless* mutant strain, which does not have trichocysts^25^. After washing them in the inorganic solution, we observed that many cells attached transiently, like the normal cells (Video S7).

Thus, *Paramecium* does not anchor on surfaces using trichocysts, nor does it stick to surfaces with its cilia. We then hypothesize that surface immobilization may be mediated by ciliary beating. We first carefully examined the posture of immobile cells in the inorganic solution (n = 21), by taking a stack of images (x20) in micrometric steps. We observed that cells immobilize in stereotypical postures. A representative example is shown in Figure 4A, observed from below the bottom surface. The cell lies on its right flank (right of the oral groove in cell frame). However, the cell’s somatic membrane is separated from the surface by a gap of about 6 µm. This gap is due to its cilia, visible at surface level (0 µm). These cilia must be tilted, since they are about 10 µm long. The membrane is in focus 6 µm above the surface, from the anterior end to about one quarter of the cell’s length from the posterior end. About 17 µm above the surface, the oral apparatus is in focus. The non-motile tail is in focus 24 µm above the surface, as well as the left edge of the oral groove. The top of the dorsal posterior region is in focus 31 µm above the surface. Thus, the cell lies along its right flank, with the oral groove on the side, facing left when observing the cell from above (Fig. 4B). This position is typical, whether cells are on the bottom or top surface (Fig. S1).

**Figure 4.**
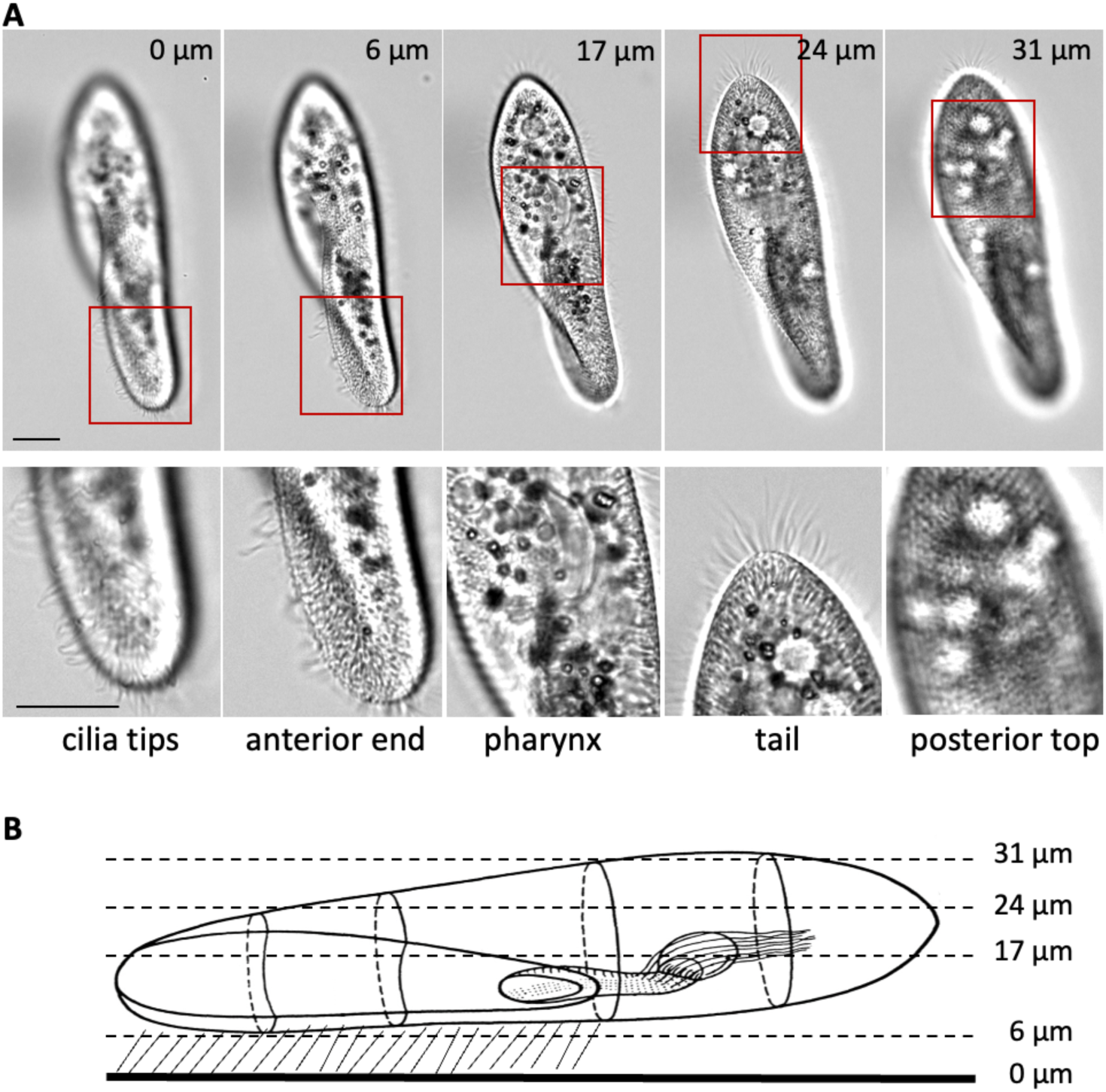
Posture of an attached cell. A, Brightfield images of a cell attached on the bottom surface, at different depths (relative to the bottom surface), using an inverted microscope. Scale bar: 20 µm. B, Inferred side view, showing the oral groove and feeding apparatus (adapted from Mast^26^).

**Figure S1.**
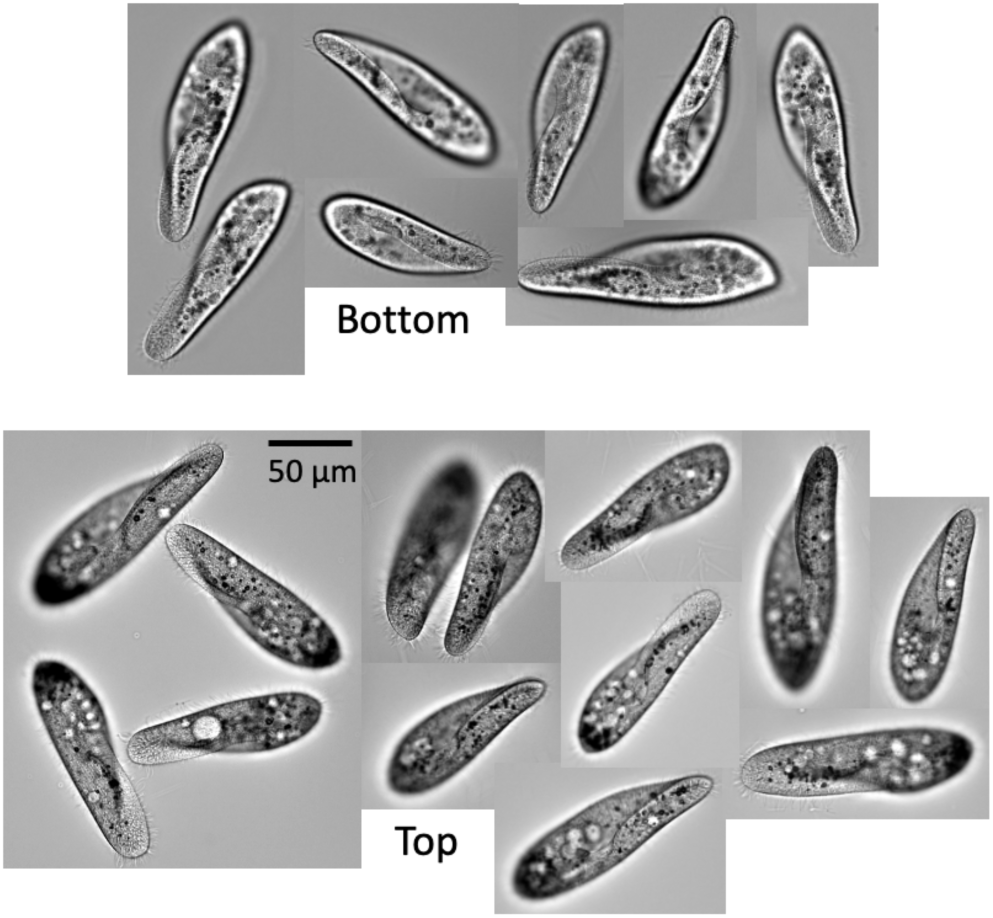
Cells attached on top and bottom (n = 21). The focus is on the anterior end, close to the surface, and cells are viewed from below (inverted microscope).

We then measured fluid motion around immobilized cells. We seeded the calcium-potassium solution with 1 µm polystyrene particles, and recorded their motion close to the surface (the focus is on the cell’s anterior end) (Fig. 5 and S2; Video S8). Figure 5A shows the flow of particles (minimum intensity projection), for a cell immobilized against the surface in the calcium-potassium solution (two other cells are shown in Figure S2A and B). Figure 5B shows the flow field and particle velocity calculated with particle image velocimetry. The flow field is strongly asymmetrical: there is a strong flow on the side of the oral groove and near the anterior end, but very little fluid motion on the dorsal side, opposite the oral groove. We compared these immobile cells with cells in the solution with added Na^+^. The latter would normally swim (Fig. 3E, F), but we make them stick to the glass slide with a bioadhesive (Cell-Tak). In contrast, in these cells, the flow field is symmetrical (Fig. 5C and D; Video S9; two other cells are shown in Figure S2C and D).

**Figure 5.**
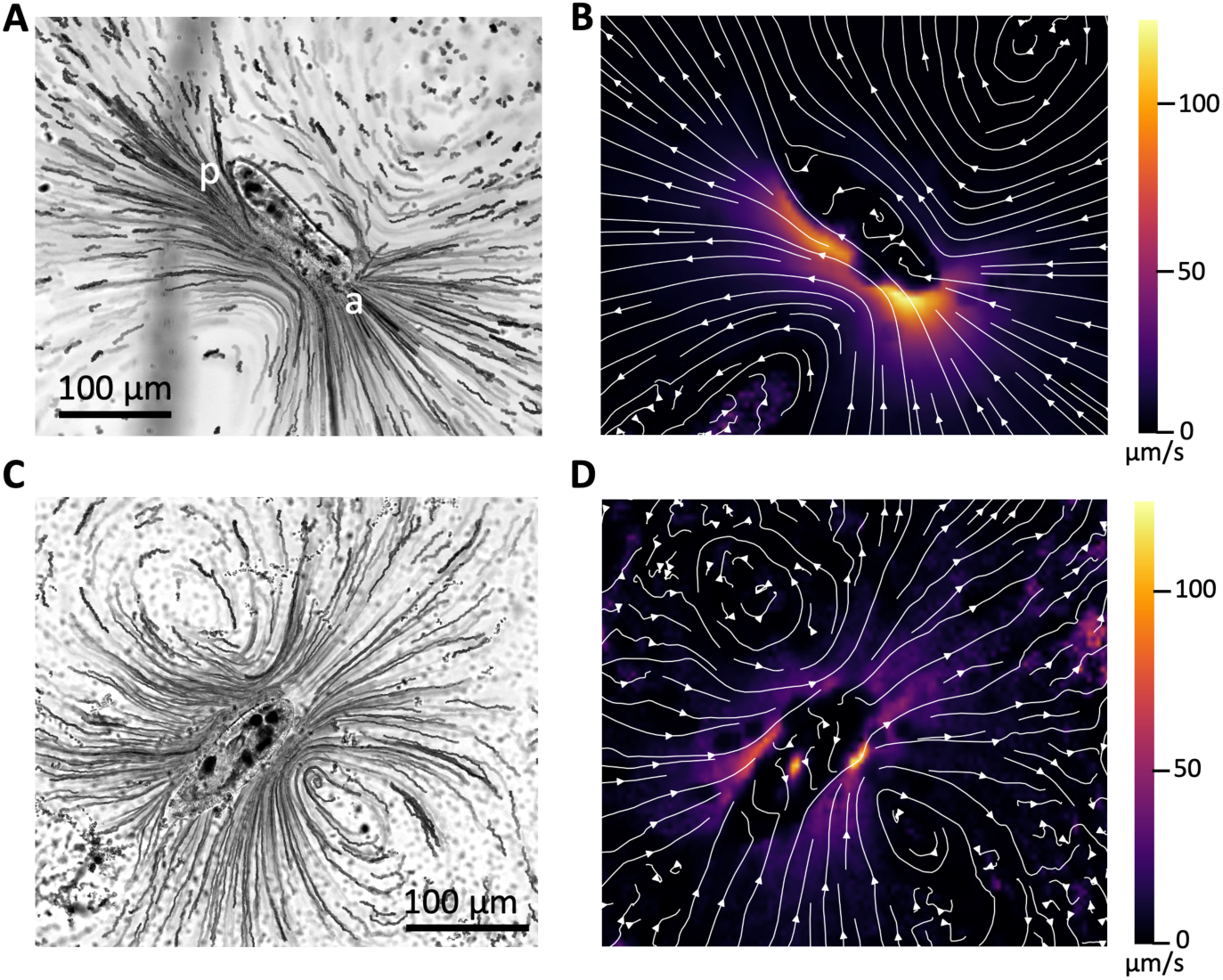
Fluid motion. A, A cell is immobilized against the bottom surface in the calcium-potassium solution seeded with 1 µm polystyrene beads, and the minimum intensity projection is shown, revealing particle motion on the side of the oral groove (view from below). The focus is on the anterior end, close to the surface (a: anterior end; p: posterior end). B, Flow field calculated by particle image velocimetry, with color-coded velocity. C, Same as A for a cell in the solution with added Na^+^ (11.5 mM), glued to the bottom surface with Cell-Tak. D, Flow field around the cell shown in C.

**Figure S2.**
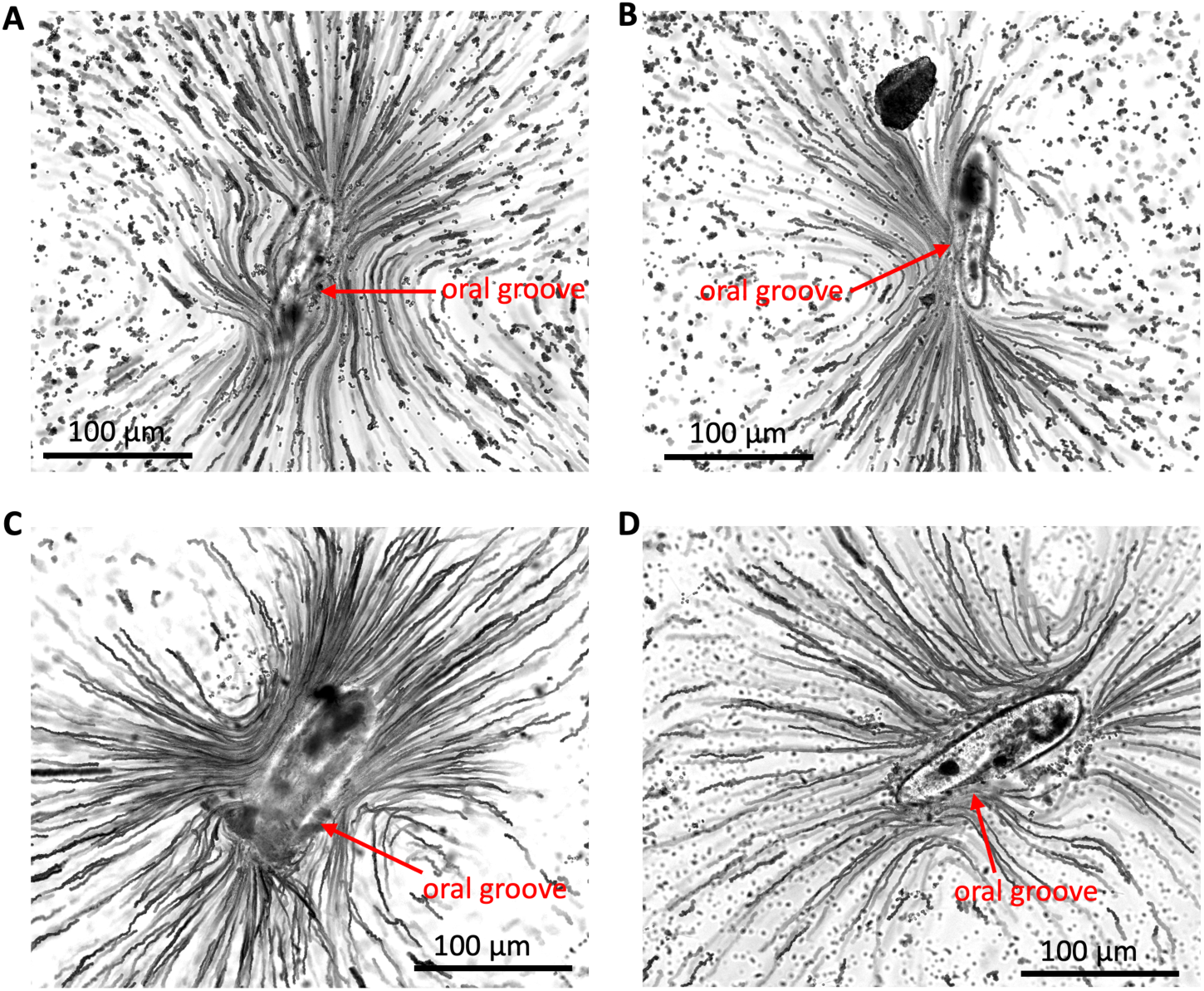
Other examples of fluid motion around cells. A, B, Two cells immobilized in the calcium-potassium solution, as in Fig. 5A. C, D, Two cells glued with Cell-Tak in a solution with extra Na^+^, as in Fig. 5C.

Thus, it appears that surface immobilization is associated with an inhibition of locomotor cilia, especially opposite the oral groove, while oral cilia continue to beat. To confirm this interpretation, we simulated the flow produced by beating cilia in a computational model, similar to previous studies^4,27^ (Fig. 6). The cell is placed near a solid wall, in a position similar to the observed position (Fig. 4), and we prescribe surface traction forces on a surface 10 µm above the cell body, in two groups: along the oral groove, and over the rest of the surface. We then vary the relative traction forces produced by oral vs. locomotor cilia. We can see that with a ratio smaller than 0.1, simulations reproduce the observed fluid motion pattern for spontaneously immobilized cells very well (Fig. 6A-C; compare with Fig. 5A, B and Fig. S2A, B). Simulations with a ratio of 0.5-1 (Fig. 6E, F) match the observed fluid motion pattern of cells attached with Cell-Tak (compare with Fig. 5C, D and Fig. S2C, D). These simulations confirm that surface immobilization corresponds to strong inhibition of locomotor cilia.

**Figure 6.**
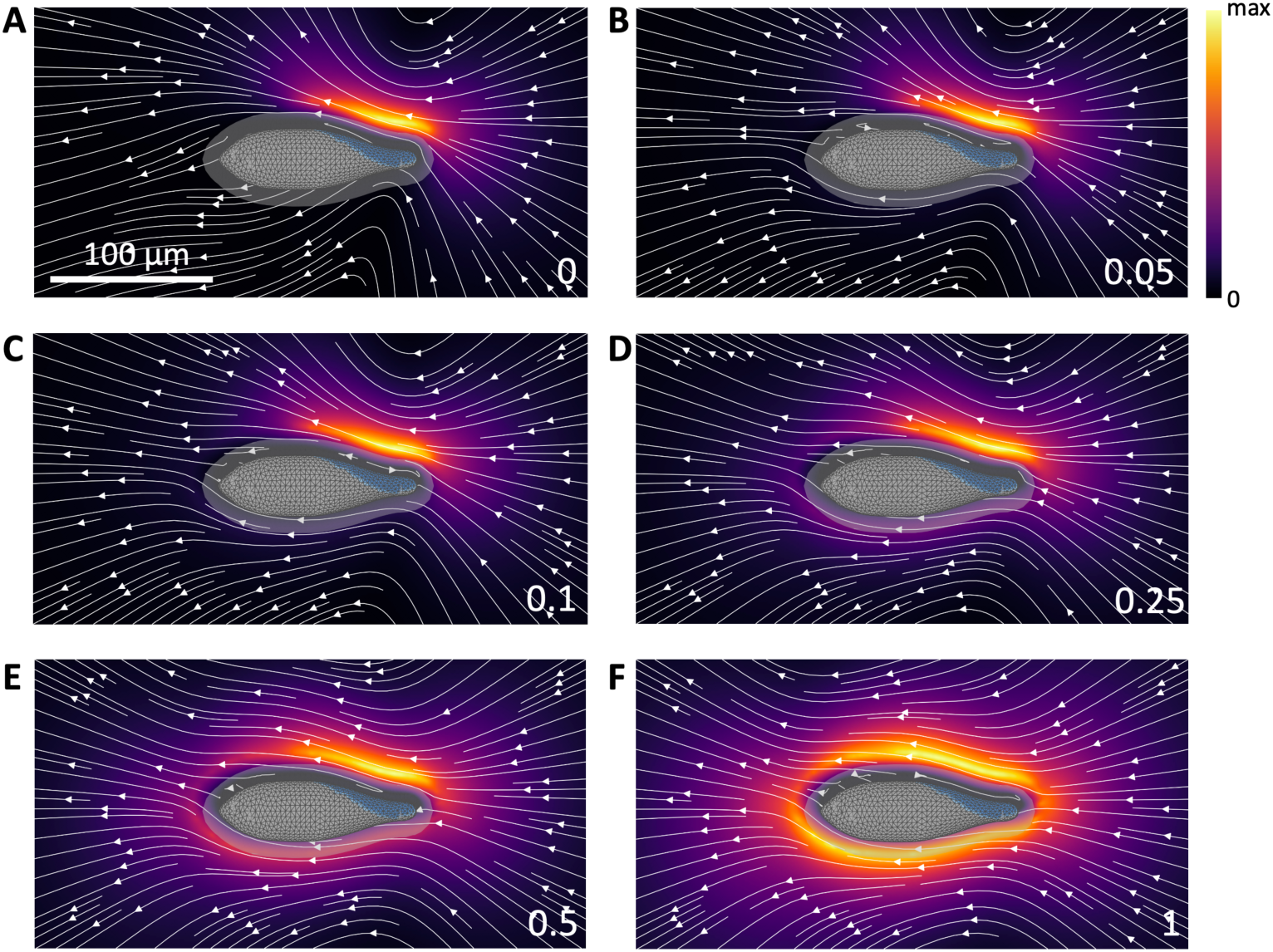
Simulation of fluid motion around a cell, seen from above. The ratio of locomotor to oral traction forces is varied from 0 to 1, and the resulting fluid velocity is shown in color (arbitrary units, normalized to the maximum velocity). The oral groove is shown in blue, and the ciliary surface is denoted by the translucent surface around the cell.

It is known that ciliary beating frequency can be regulated in *Paramecium*^28,29^. Therefore, we test whether this reduction in fluid velocity is due to a decrease in ciliary beating frequency.

#### *Paramecium* immobilizes by stopping locomotor cilia

We recorded cell images at high frame rate (500-700 Hz) to observe ciliary beating (Fig. 7). We compared immobile cells in the calcium-potassium inorganic solution with cells attached with Cell-Tak in the solution with added Na^+^ (n = 4 and n = 5, respectively). Figure 7 shows one representative cell of each group. In the calcium-potassium solution (Fig. 7A), we observe a strong beating along the oral groove, while other cilia appear to beat less strongly (Video S10). To visualize this beating, we calculated the mean difference between successive frames (Fig. 7B). On this image, the entire oral groove and oral apparatus are highlighted. The dorsal ciliary layer is visible, but weaker. Careful observation of the movies shows that many dorsal cilia are immobile, or appear to float rather than beat, as if moved passively (Video S10). To visualize immobile cilia, we calculated the temporal average of the movies at different depths, and combined parts in focus. Figure 7C shows the average images of the anterior end, focused at 6 µm above the glass surface, the middle posterior part (16 µm) and the tail (23 µm). We observe immobile cilia along the edge on the dorsal side, opposite the oral groove (the tail is always immobile). Immobile cilia are not easy to observe, since only those that stick out from the edges of the projection can be seen. Figure 7D shows another cell where the anterior end is entirely flattened, and immobile cilia are very visible (note that the oral groove appears on the other side, as the cell is against the top surface). Several authors reported that cilia touching the surface could immobilize, in particular when the cell is sliding^4,6,18^, but here the immobile cilia do not touch the surface. Therefore, it cannot be an effect of the mechanical interaction between cilia and surface, and it cannot be due to ciliary stickiness either. Rather, it appears that these cilia stop beating actively.

**Figure 7.**
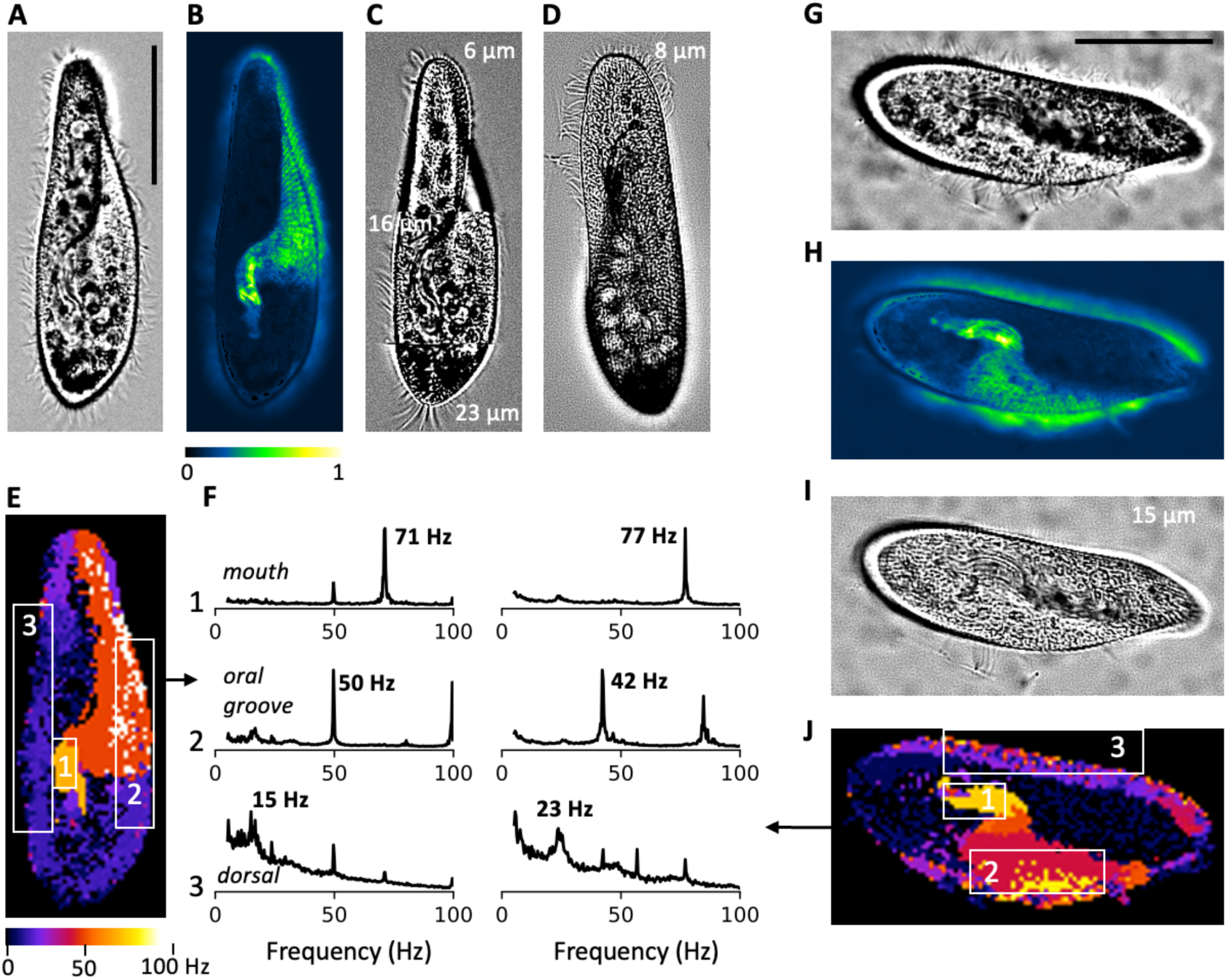
Ciliary beating. A, Cell spontaneously immobilized against the glass surface in the calcium-potassium solution. B, Mean difference between successive frames (normalized). C, Collage of mean images at three different depths relative to the glass surface, showing immobile cilia. D, Same as C, for another cell. E, Map of main beating frequency, with three highlighted regions: mouth (1), oral groove (2) and dorsal (3). F, Normalized amplitude spectrum in each of the region, showing the peak frequency. Left: cell in E; right: cell in J. G, Cell stuck to the glass surface with the bioadhesive Cell-Tak, in the solution with added Na^+^. H, Same as B in the cell shown in G. I, Mean image. A few cilia stuck to a trichocyst are visible (bottom). J, Same as E, for the cell shown in G. Scale bar: 50 µm.

We then measured the beating frequency by calculating the power spectrum across time, for each pixel. Figure 7E shows the map of peak frequency on the entire cell (same cell as in Fig. 7A-C; three other cells shown in Fig. S3). Three groups are clearly distinguished. Cilia of the mouth beat at 71 Hz, while the oral groove cilia beat at 50 Hz (Fig. 7F, left). The other cilia beat at lower frequency, with a loosely defined peak around 15 Hz. These numbers are in the range reported by previous studies^30,31^, except for the mouth, which is a new observation.

When we compare with swimming cells stuck to the glass with Cell-Tak (Fig. 7G; Video S11), the first observation is that all cilia appear to beat strongly, including dorsal cilia (Fig. 7H). When we temporally average the movie, we do not observe immobile dorsal cilia (Fig. 7I). In fact, we notice on this image a few cilia stuck to a trichocyst, on the side of the oral groove, but no immobile cilia anywhere else. Finally, when we measure beating frequency, we again observe the same three groups, with similar frequencies as in the calcium-potassium solution (Fig. 7J and 7F, right); similar observations were made in other cells (Fig. S3). Mouth beating frequency was 71±2 Hz in the calcium-potassium solution (mean ± s.d.) vs. 76±8 Hz with added Na^+^ (p=0.19, Mann-Whitney test); oral beating frequency was 52±3 Hz vs. 46±3 Hz (p=0.11, Mann-Whitney test); dorsal beating was 17±1 Hz vs. 24±2 Hz (p=0.008, Mann-Whitney one-sided test). Thus, dorsal beating frequency was slightly lower in the spontaneously attached cells, but in the range of previous reports.

**Figure S3.**
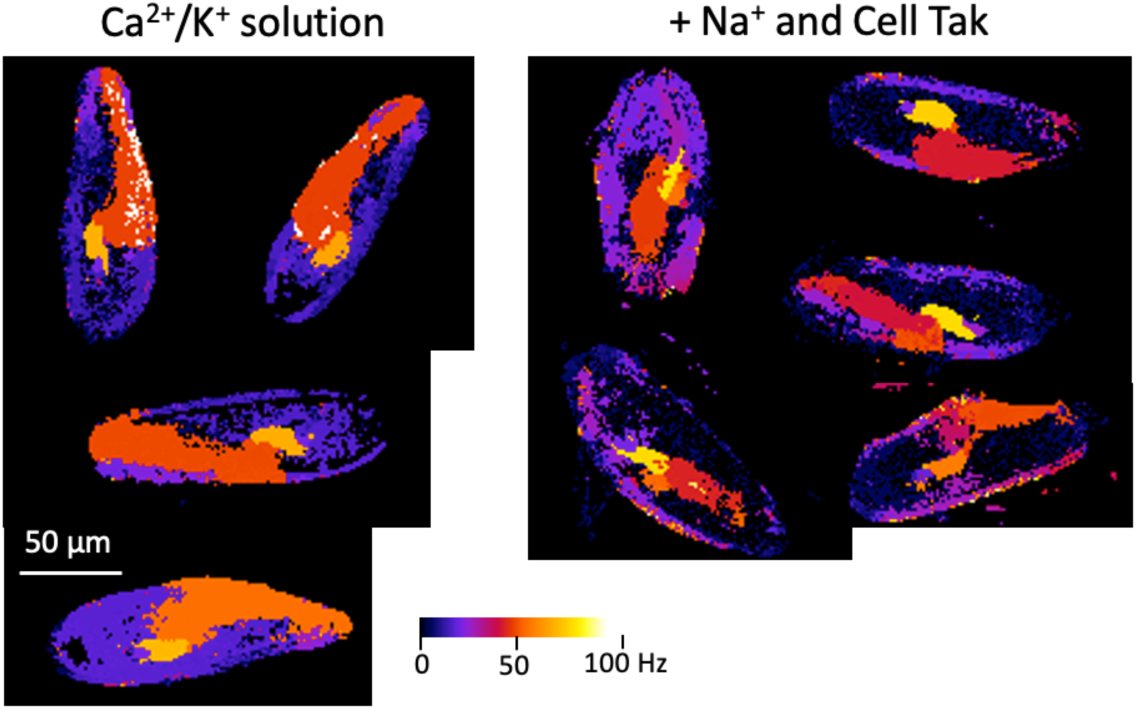
Maps of ciliary beating frequency. Left, cells spontaneously immobilized against the glass surface in the calcium-potassium solution. Right, cells stuck to the glass surface with the bioadhesive Cell-Tak, in the solution with added Na^+^.

In summary, the main salient feature of immobile cells is that many of their dorsal locomotor cilia are immobilized, either stiff or floating with no defined frequency, even though they are not in contact with the surface. We conclude that *Paramecium* immobilizes on surfaces by blocking a large part of its locomotor cilia, while oral cilia continue beating strongly.

## Discussion

We found that *Paramecium* transiently immobilizes against surfaces when it senses bacteria, by inhibiting its locomotor cilia. This is a reversible behavior in which the cell may immobilize for a few minutes before swimming away to repeat the process elsewhere. Since immobilization can occur on a surface of arbitrary orientation, the cell does not simply stop swimming and fall. Rather, we found that it immobilizes by changing its ciliary beating pattern, such that many locomotor cilia stop while oral cilia continue beating. This behavior is physiologically controlled by a circuit involving a calcium influx, which can also be artificially triggered by depletion of Na^+^ from the extracellular medium.

Our results exclude several alternative hypotheses previously proposed to explain surface attachment: general inactivity^21^ (falling), anchoring with trichocysts^19^ and increase in ciliary stickiness^16^. Several authors previously observed immobile cilia against surfaces in resting or sliding cells^4,6,18^. We confirm these observations, but we found in addition that many cilia not in contact with the surface are also immobile. This implies that cilia immobility is not directly due to the physical interaction with the surface, either mechanical (blocking the beating) or chemical (binding), but rather from physiological inhibition.

Why does *Paramecium* immobilize against surfaces? Our results connect this behavior with feeding. When bacteria are removed from the culture medium, cells do not immobilize. When immobilization is prevented by ruthenium red, growth is also inhibited. This suggests that this behavior is useful for feeding. One possibility is that *Paramecium* immobilizes where bacterial density is highest, which for immobile bacteria is probably the surfaces. Another possibility is that feeding is more efficient when the cell is in contact with a surface, for hydrodynamic reasons^32,33^, although this is disputed^34^. A physical analysis of the feeding flow in attached vs. swimming *Paramecium* could be helpful, as well as a dynamical analysis of surface attraction, extending previous work in simplified models^4,35–40^.

Our findings open a number of new questions. What physiological circuit controls this behavior? We have found that immobilization is normally triggered by the presence of bacteria, that it requires extracellular calcium, and that it is blocked by ruthenium red. This suggests that bacteria are detected by a surface receptor that opens a calcium-permeable pore, possibly of the TRP family. Since paramecia do not immobilize in the filtered medium, the detection of bacteria is not mediated by diffusible molecules secreted by bacteria, but more plausibly occurs on contact with bacteria, through either mechanical or chemical sensing.

The cell then immobilizes by selectively stopping locomotor cilia (perhaps a subset of those), which are known to have different structural and functional properties from oral cilia^30,31,41,42^. Ciliary beating is known to be modulated by several intracellular messengers, in particular calcium, cAMP and cGMP^29,43,44^, with spatially heterogeneous sensitivities^45,46^ and differentially expressed adenylate cyclases^47^. It remains an open question how these messengers can selectively control very different behaviors, such as the avoiding reaction^3^ (backward swimming followed by turning), the escape reaction^24^ (acceleration), trichocyst release^20^ and now immobilization. This study adds to the growing body of work showing complex subcellular coordination in the behavior of unicellular eukaryotes^48–51^.

More than a century ago, Jennings^1^ noted that “*from the lowest organisms up to man, behavior is essentially regulatory in character, and what we call intelligence in higher animals is a direct outgrowth of the same laws that give behavior its regulatory character in the Protozoa*”. These regulatory abilities have been demonstrated for example by studies of conditioning^2,11,12^ and adaptation^13–15^ in *Paramecium,* habituation in *Stentor*^8–10^ and *Physarum*^52^, or decision making in *Chlamydomonas*^53^. The present study highlights the striking resemblance of the feeding behavior of *Paramecium* with foraging behavior observed in animals. This is particularly intriguing, because foraging is a complex ecologically relevant behavior, involving sensory processing, memory, decision making, and motor control^54^. It also connects with a large body of theoretical literature, namely optimal foraging theory^55^. It remains to be explored to what extent *Paramecium*’s foraging behavior displays these cognitive abilities, but the perspective of studying the physiological control of foraging in a unicellular organism is appealing.

## Methods

### Bacterial cell culture

The culture medium was prepared by infusing wheatgrass powder (Pines International, Lawrence, KS, USA) in Milli-Q water, using standard protocols (see ParameciumDB wiki: https://paramecium.i2bc.paris-saclay.fr/docs/index.php/Protocols). The infusion was buffered with Tris base (6.19 mM), sodium dihydrogen phosphate monohydrate (NaH₂PO₄·H₂O, 1.45 mM), and disodium hydrogen phosphate dihydrate (Na₂HPO₄·2H₂O, 4.21 mM). The pH was adjusted to 7.0 at 22 °C using HCl. The medium was inoculated with *Klebsiella pneumoniae* (avirulent strain KpGe) and incubated at 27 °C for 16–24 h to obtain a turbid bacterial culture, then supplemented with β-sitosterol to a final concentration of 0.8 µg/ml. Bacteria are cultured every 2-3 days, then placed in the fridge.

### Paramecium culture

All experiments were carried out with the homozygous strain 51 of *P. tetraurelia*. In *Paramecium*, asexual reproduction by binary fission is associated with aging, with changes in physiology and behavior (such as sexual maturity). The clonal age is measured as the number of divisions since autogamy (self-fertilization), which can be triggered by starving in cells older than 20 divisions. To control the clonal age of cells, we designed a two-week culture protocol adapted from Beisson et al^56^. The protocol is designed to ensure vegetative growth for more than 20 divisions, so that all cells undergo autogamy when they are starved at the end of the protocol. Stationary cell density is then about 5000 cells/ml.

Cells are grown at room temperature (20-24°C) in 15 ml tubes, filled to 4 ml, with a day/night cycle (12 hours each). Every day of the week, at the same time (about 4 pm), a volume is transferred from the previous tube to a new tube, and filled to 4 ml with bacterized culture medium. The volumes are chosen so as to ensure vegetative growth with 2-3 divisions per day, as indicated in Table S1.

**Table S1.** Culture protocol, with the expected number of cell divisions and cell density.

| Week 1 |  | Monday | Tuesday | Wednesday | Thursday | Friday |
| --- | --- | --- | --- | --- | --- | --- |
|  | *Divisions* | 0 | 2-3 | 4-6 | 6-9 | 8-12 |
|  | *Density* | 5000/ml | 500-1000/ml | 350-1400/ml | 250-1950/ml | 150-2750/ml |
|  | *Picking* | 100 µl | 700 µl | 700 µl | 700 µl | 600 µl |
| *week-end at 14°C* |  |  |  |  |  |  |
| Week 2 |  | Monday | Tuesday | Wednesday | Thursday | Friday |
|  | *Divisions* | 11-15 | 13-18 | 15-21 | 17-24 | 19-27 |
|  | *Density* | 200-3300/ml | 650-1350/ml | 450-1900/ml | 300-2650/ml | 200-3700/ml |
|  | *Picking* | to 170/ml | 700 µl | 700 µl | 700 µl | 300 µl |

Cells are stationary and autogamous on the first Monday. On Friday, they are placed at 14°C, where they divide once per day. The next Monday, cell density is measured and adjusted to 170 cells/ml. On the second week-end, cells are kept at room temperature and grow to stationary phase. On the final Monday, autogamy is checked by staining them with DAPI. In case cells are not autogamous, the tube from Thursday is used.

Two cultures are maintained simultaneously with a one week shift, so as to always have young cells. The last 5 tubes of each culture line are kept. For experiments, cells are taken from first week tubes. Logarithmic cells are taken from the last tube, stationary cells are taken from a tube at least two days before.

### Cell washing

For experiments performed in media different from the original culture medium, cells were transferred through serial washing steps to minimize carryover of the original medium. A drop of cell culture and three to four drops of the experimental medium were placed on a clean Petri dish. Individual cells were sequentially transferred through the experimental medium drops using a custom-made pulled glass

Pasteur pipette, with the smallest possible volume of carryover liquid. This manipulation takes 15 to 20 minutes.

### Experimental solutions

The experimental media included a potassium-rich solution containing KCl (4 mM), CaCl₂ (1 mM), and Tris buffer (1 mM), pH 7; a sodium-rich solution with the same composition supplemented with 11.5 mM NaCl; a low-calcium solution containing KCl (4 mM), CaCl₂ (0.1 mM), and Tris buffer (1 mM); and a bacteria-free filtered culture medium prepared by passing the native culture medium of the experimental cells through a 0.22 µm filter to remove bacteria and *Paramecium* cells.

For calcium-channel inhibition experiments, ruthenium red (Sigma-Aldrich, cat. no. R2751) was added to 200 µl of cell culture medium containing cells to a final concentration of 10 µM. The suspension was gently mixed, and cells in the ruthenium red-containing medium were used directly for experimental recordings.

Similarly, EGTA (Sigma-Aldrich, St. Louis, MO, USA; cat. no. 324626) was added to 500 μl of cell culture medium containing cells to a final concentration of 0.32 μM and 0.38 μM. The suspension was gently mixed, and cells from that suspension were used directly for experimental recordings.

### Observation chambers

Experimental droplets (30–50 µl) were placed in two types of observation chambers (Fig. S4). In the first setup, the droplet was sandwiched between the outer surface of an inverted 60 mm Petri dish lid and the bottom of a larger Petri dish (150 mm). The rim of the inverted lid provided a spacer to maintain a separation of about 880 µm between the two surfaces. The surrounding space was humidified with moistened tissue to minimize evaporation, and the larger Petri dish was covered with its lid to create a closed, humid, and transparent observation chamber. This configuration slattened the droplet, providing improved optical access and reducing distortion caused by the curved meniscus of an open drop.

**Figure S4.**
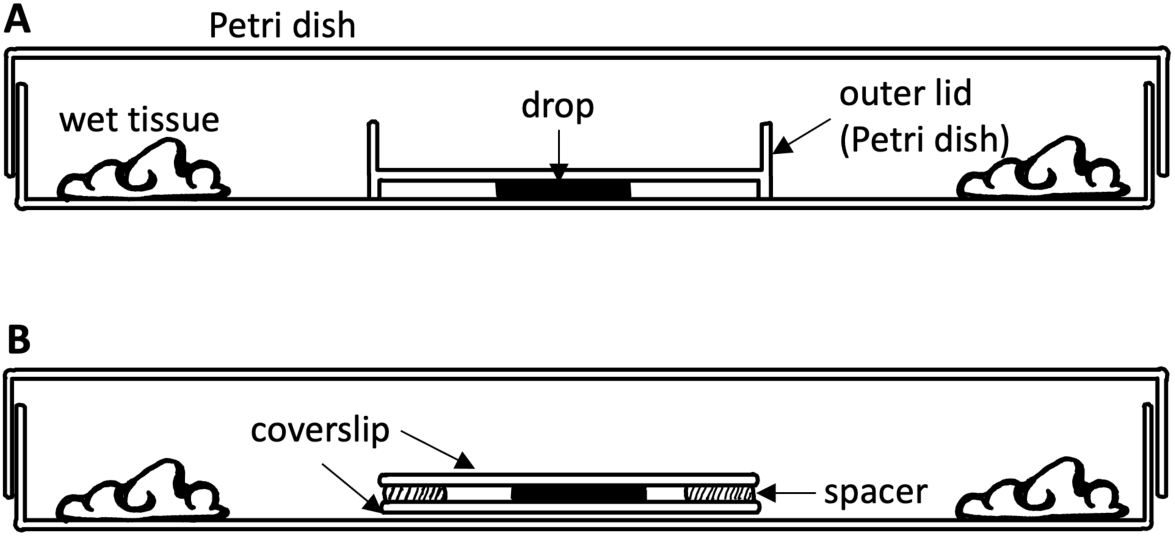
Observation chambers. A, Drop on Petri dish. B, Drop between glass coverslips.

In the second setup, experimental droplets were confined between two glass coverslips separated by a spacer of about 500 µm and maintained inside a larger Petri dish (150 mm) with moistened tissue to minimize evaporation.

### Immobilization

To immobilize cells, we coated glass coverslips with Cell-Tak (Sigma-Aldrich, cat. no. CLS354240), with a protocol adapted from Bell et al^30^. The coating solution was prepared by diluting Cell-Tak in 10 mM Tris buffer (pH 8.0) and activating it with 1 M NaOH added at a volume equal to half of the Cell-Tak volume. The volume of coating solution applied to each coverslip was adjusted to achieve a final Cell-Tak coating density of 10 μg cm⁻². After application, coverslips were incubated at room temperature overnight in an enclosed chamber and then rinsed thoroughly with deionized water.

### Image acquisition

Population dynamics were imaged using a stereomicroscope (SMZ800N, Nikon) equipped with a Plan Apo 0.5× objective (WD 82 mm, Nikon) and a 16 MP monochrome camera (ace 2 a2A5320-23umBAS, Basler). Pixel size was 5.06 µm, and the field of view was 26.9 × 15.35 mm. Image frames were acquired at a rate of 20 Hz and processed in real time to track cells (see *Cell tracking* below).

Cells in Fig. 1B are shown with an inverted microscope (Nikon Ti2-U) with 40× objective. High-magnification imaging for particle image velocimetry (PIV) and ciliary beating frequency measurements was performed using an inverted microscope (Axio Observer, ZEISS) with transmitted LED illumination, a 20× Plan Apochromat air objective (NA 0.8, ZEISS), and a monochrome camera (Axiocam 807 mono, ZEISS). Images were acquired at 92-245 Hz for PIV measurements, and at 500-700 Hz for ciliary beating measurements.

### Cell tracking

Images were acquired at 20 Hz and stored as raw tiff files. Files were then processed online to localize and track cells as follows. The background is calculated as the mean frame over a 10-30 s interval, and recalculated on subsequent intervals. It is then subtracted from the current frame and the resulting image is thresholded then saved with lossless compression. The background image is also stored. This ensures manageable data size for storage. To extract cell positions, we binarize the thresholded image, select connected components of adequate size, and calculate the centroid, length and width. Cell positions are then connected over frames into tracks using the python package *norfair*^57^, which is based on a Kalman filter. When raw images have been processed, they are deleted. This allows us to process and save very long video recordings (up to 12 hours; in full field, this represents 14 TB of raw images).

Immobile cells appear in the background images. To identify them, we use the same software as for tracking cells, but applied to the sequence of backgrounds, with the mean image subtracted.

### Counting cells

Cells in each drop are selected by manually defining a rectangular region of interest, or by segmenting the drop with edge detection. We then select cells with plausible dimensions, to reduce identification errors. The median length *l* and width *w* are calculated over the entire recording, then we select cells with length in [*l*/2, 3*l*/2] and width in [*w*/2, 2*w*]. It is more robust to use median values than fixed dimensions, because the apparent dimensions can depend on the binarization threshold.

In each frame, we count the number of swimming cells, defined as those with an instantaneous speed greater than 30 µm/s. We then calculate the maximum number of swimming cells in each background interval. In this way, directional changes or transient immobilization do not bias the estimate. This number is then compared to the number of cells in the background recording. For visualization, the results are smoothed with a 5 min rolling window. In Fig. 1H, immobile cells cannot be counted in the background, because cells form aggregates. Instead, we show the number of swimming cells relative to the initial number of cells (this goes slightly above 100% because of a few cell divisions).

### Attachment duration

To calculate attachment duration (Fig. 1F), we extract immobile tracks from the background images. Using *trackpy*, identified cells are connected over successive background images only if they are less than 50 µm away, and allowing the cell to disappear for up to 5 images.

### Statistics

Statistics of immobilization are shown in Fig. 2B, 2D, 3D and 3F. The numbers of swimming and immobile cells are smoothed with a 5 min rolling window. We then calculate the maximum proportion of immobile cells. The error bar is the standard error of the mean, calculated from a binomial distribution with parameters *p* the maximum proportion of immobile cells and *n* the total number of cells at that time (i.e., assuming cells are independent). Statistical significance is calculated with Welch’s t-test applied to binomial variables.

### Ciliary beating

Immobile cells are imaged at 500-700 Hz for 30 s. We then extract the 2000 frames interval where the cell is most stable. The background (mean image) is subtracted, thresholded, the cell centroid is calculated in each frame, and then we select the interval with smallest centroid variance. We then calculate the power spectrum (across time) of each pixel, using a Fourier transform. In Fig. 7F, this spectrum is averaged over a region of interest, and visualized between 5 Hz and 100 Hz. In frequency maps (Fig. 7E, 7J), we bin the power spectrum in squares of 3×3 pixels, smooth it with a low pass filter (uniform with a size 3 frequency bins), then select the peak frequency between 5 Hz and 100 Hz in each bin. Finally, bins where total power is smaller than 1% of the maximum total power (across space) are masked (shown in black), as those where the peak frequency is on the ends of the frequency interval (i.e., 5 and 100 Hz).

### Particle image velocimetry

To measure the slow of particles around cells, we used particle image velocimetry (PIV). We seeded the experimental medium with 1 μm polystyrene microspheres (Sigma-Aldrich, product no. 89904) at a 1:2500 dilution of the stock suspension, corresponding to 7.6 10^7^ particles/ml. Subsequently, 10–15 μl of cells suspended in experimental medium containing tracer particles was transferred onto the glass surface.

Video recordings (frame rate 92-245 Hz) are pre-processed by subtracting the background (mean image), and band-pass filtering to enhance particle visibility, using a difference of Gaussians with standard deviations *σ* = 1 µm (particle size) and 1.3*σ*. We then used the PIV package *openpiv* with 10 µm analysis windows.

### Hydrodynamic simulation

To calculate the sluid slow around the cell expected from a given pattern of ciliary beating, we model the cell body as a rigid surface with zero velocity, representing a cell attached to or stationary near a solid wall. This computational framework was first introduced by Ohmura et al.^4^ and further analyzed by Ishikawa et al^27^. The slow is driven by a layer of prescribed surface traction forces located 10 µm above the cell body, representing the time-averaged force exerted by the beating cilia. The traction forces for the locomotor cilia are directed posteriorly and to the right, while those for the oral groove are directed along the oral groove toward the cell mouth. We vary the relative strengths of these two types of ciliary forces to examine how changes in locomotor ciliary activity alter the resulting slow (Fig. 6). The cell is positioned a ciliary width, or 10 µm, above the surface and oriented so that it rests on its right slank, as in Figure 4. The resulting slow fields are computed using the method of images for regularized Stokeslets^58^. Additional details of the numerical implementation are provided in the Supplementary Methods.

## Code and data availability

Code for figures and data analysis, including hydrodynamics simulations, can be found at https://github.com/romainbrette/Paramecium-thigmotaxis. Code for cell tracking can be found at https://github.com/mstimberg/online_bg_removal. Data can be found at https://zenodo.org/uploads/21509306.

## Acknowledgments

We thank Mireille Bétermier and Linda Sperling for providing mutant strains of *Paramecium*.

This work was supported by Agence Nationale de la Recherche (ANR-20-CE30-0025-01, ANR-21-CE16-0013-02, ANR-23-CE16-0020-02, ROBOTEX ANR-10-EQPX-44-01 and TIRREX ANR-21-ESRE-0015) and by the National Science Foundation (CBET-2516633).

## References

1. Jennings (1906). Behavior of the lower organisms (New York, The Columbia university press, The Macmillan company, agents; [etc., etc.]).

2. Brette, R. (2021). Integrative Neuroscience of Paramecium, a “Swimming Neuron.” eNeuro 8, ENEURO.0018-21.2021. 10.1523/ENEURO.0018-21.2021.

3. Eckert, R. (1972). Bioelectric Control of Ciliary Activity. Science 176, 473–481. 10.1126/science.176.4034.473.

4. Ohmura, T., Nishigami, Y., Taniguchi, A., Nonaka, S., Manabe, J., Ishikawa, T., and Ichikawa, M. (2018). Simple mechanosense and response of cilia motion reveal the intrinsic habits of ciliates. Proc. Natl. Acad. Sci. 115, 3231–3236. 10.1073/pnas.1718294115.

5. Escoubet, N., Brette, R., Pontani, L.-L., and Prevost, A.M. (2023). Interaction of the mechanosensitive microswimmer Paramecium with obstacles. R. Soc. Open Sci. 10, 221645. 10.1098/rsos.221645.

6. Jennings, H.S. (1897). Studies on Reactions to Stimuli in Unicellular Organisms. J. Physiol. 21, 258–322.

7. Dexter, J.P., Prabakaran, S., and Gunawardena, J. (2019). A Complex Hierarchy of Avoidance Behaviors in a Single-Cell Eukaryote. Curr. Biol. CB 29, 4323–4329.e2. 10.1016/j.cub.2019.10.059.

8. Rajan, D., Makushok, T., Kalish, A., Acuna, L., Bonville, A., Correa Almanza, K., Garibay, B., Tang, E., Voss, M., Lin, A., et al. (2023). Single-cell analysis of habituation in Stentor coeruleus. Curr. Biol. CB 33, 241–251.e4. 10.1016/j.cub.2022.11.010.

9. Wood, D.C. (1969). Parametric studies of the response decrement produced by mechanical stimuli in the protozoan, Stentor coeruleus. J. Neurobiol. 1, 345–360. 10.1002/neu.480010309.

10. Rajan, D.H., Albright, A., Kim, H., Diaz, U., Hudnall, Y., Steube, N., Dey, G., Liu, T., and Marshall, W.F. (2026). Molecular pathways for learning in the single-cell Stentor coeruleus. Curr. Biol. 36, 2367–2381.e9. 10.1016/j.cub.2026.03.080.

11. Hennessey, T.M., Rucker, W.B., and McDiarmid, C.G. (1979). Classical conditioning in paramecia. Anim. Learn. Behav. 7, 417–423. 10.3758/BF03209695.

12. Gershman, S.J., Balbi, P.E., Gallistel, C.R., and Gunawardena, J. (2021). Reconsidering the evidence for learning in single cells. eLife 10, e61907. 10.7554/eLife.61907.

13. Nakaoka, Y., Tokui, H., Gion, Y., Inoue, S., and Oosawa, F. (1982). Behavioral Adaptation of Paramecium caudatum to Environmental Temperature. Proc. Jpn. Acad. Ser. B 58, 213–217. 10.2183/pjab.58.213.

14. Oka, T., Nakaoka, Y., and Oosawa, F. (1986). Changes in Membrane Potential during Adaptation to External Potassium Ions in Paramecium Caudatum. J. Exp. Biol. 126, 111–117.

15. Preston, R.R., and Hammond, J.A. (1998). Long-term adaptation of Ca2+-dependent behaviour in Paramecium tetraurelia. J. Exp. Biol. 201, 1835–1846.

16. Kitamura, A. (1982). Attachment of Paramecium to polystyrene surfaces: a model system for the analysis of sexual cell recognition and nuclear activation. J. Cell Sci. 58, 185–199. 10.1242/jcs.58.1.185.

17. Iwatsuki, K., and Hirano, T. (1996). An increase in the influx of calcium ions into cilia induces thigmotaxis inParamecium caudatum. Experientia 52, 831–833. 10.1007/BF01923998.

18. Iwatsuki, K., and Hirano, T. (1995). Induction of the thigmotaxis in Paramecium caudatum. Comp. Biochem. Physiol. A Physiol. 110, 167–170. 10.1016/0300-9629(94)00125-D.

19. Saunders, J.T. (1925). The Trichocysts of Paramecium. Biol. Rev. 1, 249–269. 10.1111/j.1469-185X.1925.tb00554.x.

20. Plattner, H. (2017). Trichocysts-Paramecium’s Projectile-like Secretory Organelles: Reappraisal of their Biogenesis, Composition, Intracellular Transport, and Possible Functions. J. Eukaryot. Microbiol. 64, 106–133. 10.1111/jeu.12332.

21. Machemer, H., Machemer-Röhnisch, S., and Bräucker, R. (1993). Velocity and Graviresponses in Paramecium during Adaptation and Varied Oxygen Concentrations. Arch. Für Protistenkd. 143, 285–296. 10.1016/S0003-9365(11)80325-8.

22. Machemer, H. (1996). A theory of gravikinesis in Paramecium. Adv. Space Res. 17, 11–20. 10.1016/0273-1177(95)00607-G.

23. Lodh, S., Yano, J., Valentine, M.S., and Van Houten, J.L. (2016). Voltage-gated calcium channels of Paramecium cilia. J. Exp. Biol. 219, 3028–3038. 10.1242/jeb.141234.

24. Nakaoka, Y., and Iwatsuki, K. (1992). Hyperpolarization-activated inward current associated with the frequency increase in ciliary beating of Paramecium. J. Comp. Physiol. A 170, 723–727. 10.1007/BF00198983.

25. Pollack, S. (1974). Mutations affecting the trichocysts in Paramecium aurelia. I. Morphology and description of the mutants. J. Protozool. 21, 352–362. 10.1111/j.1550-7408.1974.tb03669.x.

26. Mast, S.O. (1947). The food-vacuole in paramecium. Biol. Bull. 92, 31–72. 10.2307/1537967.

27. Ishikawa, T., Pedley, T.J., Drescher, K., and Goldstein, R.E. (2020). Stability of dancing Volvox. J. Fluid Mech. 903, A11. 10.1017/jfm.2020.613.

28. Machemer, D.H., and Eckert, D.R. (1975). Ciliary frequency and orientational responses to clamped voltage steps inParamecium. J. Comp. Physiol. 104, 247–260. 10.1007/BF01379051.

29. Nakaoka, Y., Tanaka, H., and Oosawa, F. (1984). Ca2+-dependent regulation of beat frequency of cilia in Paramecium. J. Cell Sci. 65, 223–231.

30. Bell, W.E., Hallworth, R., Wyatt, T.A., and Sisson, J.H. (2015). Use of a Novel Cell Adhesion Method and Digital Measurement to Show Stimulus-dependent Variation in Somatic and Oral Ciliary Beat Frequency in Paramecium. J. Eukaryot. Microbiol. 62, 144–148. 10.1111/jeu.12153.

31. Laan, D.M., Kourkoulou, A.M., and Ramirez-San-Juan, G.R. (2026). Heterogeneity in cilia patterning enables multiple flow functions within a single cell. Preprint at bioRxiv, 10.64898/2026.02.19.706812 https://doi.org/10.64898/2026.02.19.706812.

32. Jonsson, P.R., Johansson, M., and Pierce, R.W. (2004). Attachment to suspended particles may improve foraging and reduce predation risk for tintinnid ciliates. Limnol. Oceanogr. 49, 1907–1914. 10.4319/lo.2004.49.6.1907.

33. Christensen-Dalsgaard, K.K., and Fenchel, T. (2003). Increased filtration efficiency of attached compared to free-swimming flagellates. Aquat. Microb. Ecol. 33, 77–86. 10.3354/ame033077.

34. Liu, J., Man, Y., Costello, J.H., and Kanso, E. (2026). Feeding rates in sessile versus motile ciliates are hydrodynamically equivalent. eLife 13, RP99003. 10.7554/eLife.99003.

35. Berke, A.P., Turner, L., Berg, H.C., and Lauga, E. (2008). Hydrodynamic attraction of swimming microorganisms by surfaces. Phys. Rev. Lett. 101, 038102. 10.1103/PhysRevLett.101.038102.

36. Ishimoto, K., and Gaffney, E.A. (2013). Squirmer dynamics near a boundary. Phys. Rev. E 88, 062702. 10.1103/PhysRevE.88.062702.

37. Li, G.-J., and Ardekani, A.M. (2014). Hydrodynamic interaction of micro-swimmers near a wall. Phys. Rev. E Stat. Nonlin. Soft Matter Phys. 90, 013010.

38. Lintuvuori, J.S., Brown, A.T., Stratford, K., and Marenduzzo, D. (2016). Hydrodynamic oscillations and variable swimming speed in squirmers close to repulsive walls. Soft Matter 12, 7959–7968. 10.1039/C6SM01353H.

39. Spagnolie, S.E., and Lauga, E. (2012). Hydrodynamics of self-propulsion near a boundary: predictions and accuracy of far-field approximations. J. Fluid Mech. 700, 105–147. 10.1017/jfm.2012.101.

40. Echigoya, S., Ohmura, T., Sato, K., Nakagaki, T., and Nishigami, Y. (2026). Geometrical preference of anchoring sites in the unicellular organism Stentor coeruleus. Proc. Natl. Acad. Sci. 123, e2518816123. 10.1073/pnas.2518816123.

41. Funfak, A., Fisch, C., Abdel Motaal, H.T., Diener, J., Combettes, L., Baroud, C.N., and Dupuis-Williams, P. (2015). Paramecium swimming and ciliary beating patterns: a study on four RNA interference mutations. Integr. Biol. Quant. Biosci. Nano Macro 7, 90–100. 10.1039/c4ib00181h.

42. Aubusson-Fleury, A., Cohen, J., and Lemullois, M. (2015). Ciliary heterogeneity within a single cell: the Paramecium model. Methods Cell Biol. 127, 457–485. 10.1016/bs.mcb.2014.12.007.

43. Bonini, N.M., and Nelson, D.L. (1988). Differential regulation of Paramecium ciliary motility by cAMP and cGMP. J. Cell Biol. 106, 1615–1623. 10.1083/jcb.106.5.1615.

44. Walczak, C.E., and Nelson, D.L. (1994). Regulation of dynein-driven motility in cilia and flagella. Cell Motil. 27, 101–107. 10.1002/cm.970270202.

45. Noguchi, M., Nakamura, Y., and Okamoto, K.-I. (1991). Control of ciliary orientation in ciliated sheets from Paramecium–differential distribution of sensitivity to cyclic nucleotides. Cell Motil. 20, 38–46. 10.1002/cm.970200105.

46. Noguchi, M., Kurahashi, S., Kamachi, H., and Inoue, H. (2004). Control of the Ciliary Beat by Cyclic Nucleotides in Intact Cortical Sheets from Paramecium. Zoolog. Sci. 21, 1167–1175. 10.2108/zsj.21.1167.

47. Kawano, M., Tominaga, T., Ishida, M., and Hori, M. (2020). Roles of Adenylate Cyclases in Ciliary Responses of Paramecium to Mechanical Stimulation. J. Eukaryot. Microbiol. 67, 532–540. 10.1111/jeu.12800.

48. Larson, B.T., Garbus, J., Pollack, J.B., and Marshall, W.F. (2022). A unicellular walker controlled by a microtubule-based finite-state machine. Curr. Biol. CB 32, 3745–3757.e7. 10.1016/j.cub.2022.07.034.

49. Wan, K.Y. (2018). Coordination of eukaryotic cilia and flagella. Essays Biochem. 62, 829–838. 10.1042/EBC20180029.

50. Wan, K.Y., and Goldstein, R.E. (2017). Run stop shock, run shock run: Spontaneous and stimulated gait-switching in a unicellular octoflagellate. ArXiv170607922 Cond-Mat Q-Bio.

51. Laeverenz-Schlogelhofer, H., and Wan, K.Y. (2024). Bioelectric control of locomotor gaits in the walking ciliate Euplotes. Curr. Biol. 34, 697–709.e6. 10.1016/j.cub.2023.12.051.

52. Boisseau, R.P., Vogel, D., and Dussutour, A. (2016). Habituation in non-neural organisms: evidence from slime moulds. Proc. R. Soc. B Biol. Sci. 283, 20160446. 10.1098/rspb.2016.0446.

53. Raikwar, S., Al-Kassem, A., Gov, N.S., Pesci, A.I., Jeanneret, R., and Goldstein, R.E. (2025). Phototactic Decision-Making by Microalgae. Phys. Rev. Lett. 135, 228401. 10.1103/3fry-7tsw.

54. Dennis, E.J., and Hady, A.E. (2026). Neurobiology of Foraging: An Integrative Approach. 10.1146/annurev-neuro-091724-040841.

55. Grima, L.L., Haberkern, H., Mohanta, R., Morimoto, M.M., Rajagopalan, A.E., and Scholey, E.V. (2025). Foraging as an ethological framework for neuroscience. Trends Neurosci. 10.1016/j.tins.2025.08.006.

56. Beisson, J., Bétermier, M., Bré, M.-H., Cohen, J., Duharcourt, S., Duret, L., Kung, C., Malinsky, S., Meyer, E., Preer, J.R., et al. (2010). Maintaining Clonal Paramecium tetraurelia Cell Lines of Controlled Age through Daily Reisolation. Cold Spring Harb. Protoc. 2010, pdb.prot5361. 10.1101/pdb.prot5361.

57. Alori, J., Descoins, A., javier, Lezama, F., KotaYuhara, Fernández, D., Castro, A., fatih, David, Linares, R.C., et al. (2023). tryolabs/norfair: v2.2.0. 10.5281/zenodo.7504727.

58. Ainley, J., Durkin, S., Embid, R., Boindala, P., and Cortez, R. (2008). The method of images for regularized Stokeslets. J. Comput. Phys. 227, 4600–4616. 10.1016/j.jcp.2008.01.032.

