## Supplementary Methods for "Active control of surface immobilization in the foraging behavior of *Paramecium*"

### Supplementary Information for Active control of surface immobilization in the foraging behavior of *Paramecium*

#### Overview

We represent the cell surface using a parameterized description of its geometry. To construct the cell body, we first mesh an ellipsoid using *DistMesh* [3], with an aspect ratio matching that of an image of the *Paramecium tetraurelia* used in this study. The mesh points are projected onto an axisymmetric surface that defines the cell body surface. A second set of points, representing the ciliary traction layer, is generated by extending these mesh points outward from the cell body surface. The traction layer accounts for two populations of cilia, each with a different strength and direction. With traction forces prescribed, we use the method of images for regularized Stokeslets (MIRS) [1] to determine the forces required on the cell body to satisfy the no-slip boundary condition. Finally, the forces on the cell body and traction layer are used to calculate the surrounding fluid velocity.

#### 1 Surface Parametrization

*Paramecia* are commonly modeled as being azimuthally symmetric about their main, or anteroposterior, axis. We therefore use a cylindrical coordinate system $\{\rho, z\}$, parameterized by the angular coordinates $\{\theta, \varphi\}$. Here, $\theta$ is the polar angle, with $\theta = 0$ at the anterior and $\theta = \pi$ at the posterior, while $\varphi$ is the azimuthal angle, with $\varphi = 0$ corresponding to the cell mouth. Based on an image of *Paramecium tetraurelia* used in this study, a set of points on the boundary is manually selected, and a piecewise cubic Hermite interpolating polynomial is constructed through those points to represent the profile functions $\rho(\theta)$ and $z(\theta)$.

Figure 1 illustrates the cylindrical coordinates and local orthonormal basis used to describe the cell body surface. We parameterize this surface in Cartesian coordinates as

$$\mathbf{X}(\theta, \varphi) = \Big(\rho(\theta)\cos\varphi,\ \rho(\theta)\sin\varphi,\ z(\theta)\Big). \tag{1}$$

We compute an orthonormal basis $\{\hat{\theta}, \hat{\varphi}, \hat{n}\}$ on this surface, with the explicit $\theta$-dependence of $\rho$ and $z$ surpressed for brevity. The tangent vector in the $\theta$ direction is

$$\frac{\partial \mathbf{X}}{\partial \theta} = \Big(\rho'\cos\varphi,\ \rho'\sin\varphi,\ z'\Big), \tag{2}$$

giving the normalized tangent vector

$$\hat{\theta} = \frac{1}{\sqrt{(\rho')^2 + (z')^2}}\Big(\rho'\cos\varphi,\ \rho'\sin\varphi,\ z'\Big). \tag{3}$$

Similarly, the tangent vector in the $\varphi$ direction is

$$\hat{\varphi} = \Big(-\sin\varphi,\ \cos\varphi,\ 0\Big). \tag{4}$$

The outward unit normal is then

$$\hat{n} = \hat{\theta} \times \hat{\varphi} = \frac{1}{\sqrt{(\rho')^2 + (z')^2}}\Big(-z'\cos\varphi,\ -z'\sin\varphi,\ \rho'\Big). \tag{5}$$

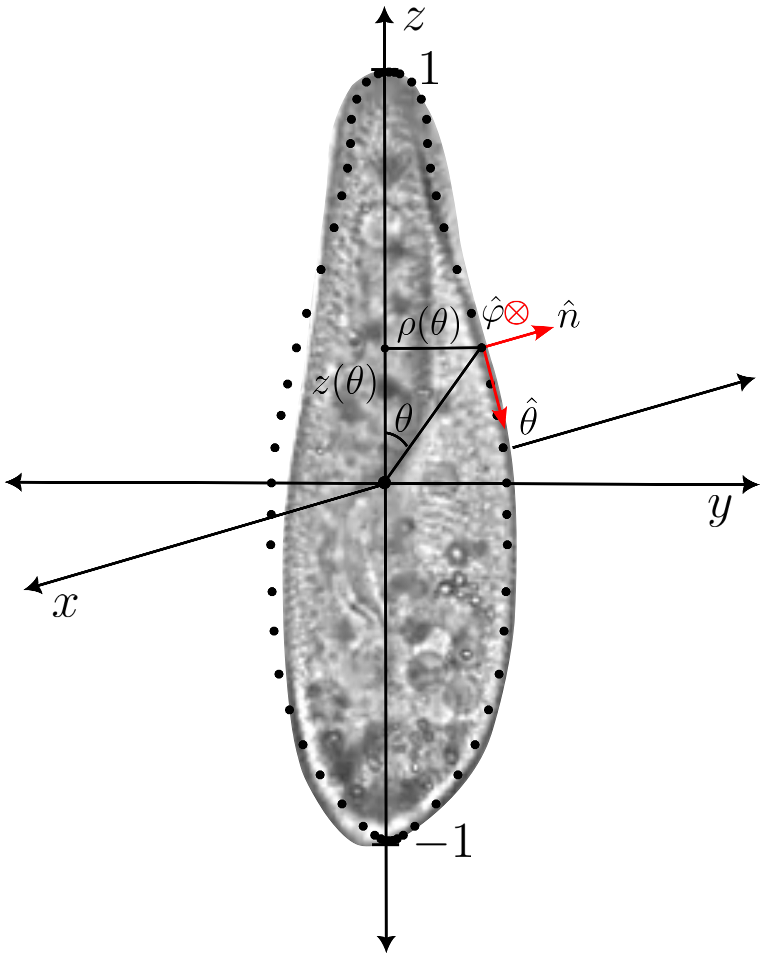


Figure 1: Coordinate system used to parameterize the *Paramecium tetraurelia* cell body surface. The manually selected points used to construct the interpolated cell profile are shown together with the cylindrical coordinates $\rho$ and $z$, which describe the position of a point on the axisymmetric surface. The set $\{\hat{\theta}, \hat{\varphi}, \hat{n}\}$ denotes the local orthonormal basis, consisting of the polar, azimuthal, and outward normal directions, respectively. The cell is nondimensionalized such that the anterior and posterior ends are located at $z = 1$ and $z = -1$, respectively, corresponding to a total representative cell length of 120 µm.

#### 2 Surface Meshing and the Traction Layer

The mesh on an ellipsoid described above is mapped onto the parameterized cell body surface by selectively inwardly- or outwardly-projecting points in the $\rho$-direction, while keeping their $z$-coordinates fixed. Figure 2 shows the mesh before and after this projection.

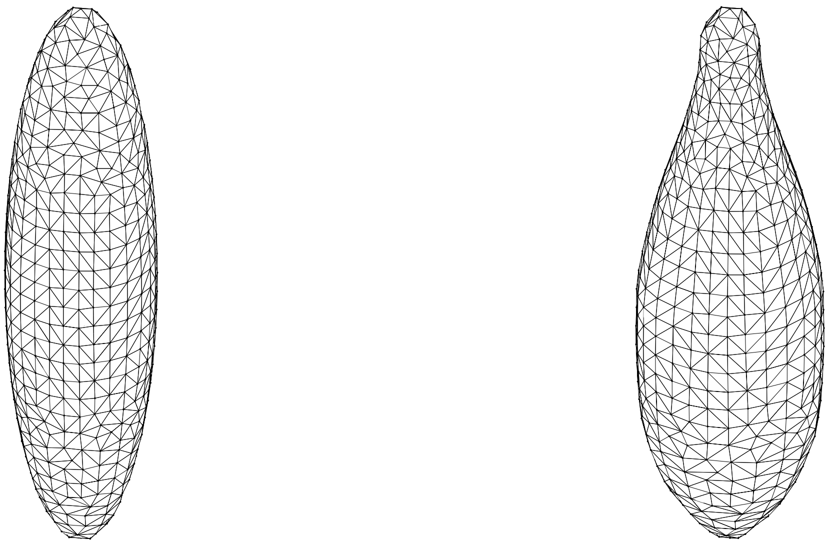

Figure 2: Construction of the cell body surface mesh. (Left) Meshed ellipsoid generated using *DistMesh* [3]. (Right) The same mesh after projection in the $\rho$-direction onto the parameterized cell surface.

The resulting cell body surface is represented by the set of points

$$X_b = \{\mathbf{x}_{b,i}\}_{i=1}^{N}. \tag{6}$$

Each body point $\mathbf{x}_{b,i}$ is associated with angular coordinates $(\theta_i, \varphi_i)$ and the corresponding orthonormal basis $\{\hat{\theta}_i, \hat{\varphi}_i, \hat{n}_i\}$ defined in the previous section. The traction layer is constructed by projecting each cell body point outward along its surface normal by a fixed distance $\epsilon$, representing the effective width of the ciliary layer. The traction-layer points are therefore given by

$$X_t = \{\mathbf{x}_{t,i} \mid \mathbf{x}_{t,i} = \mathbf{x}_{b,i} + \epsilon \hat{n}_i\}\,. \tag{7}$$

Each traction-layer point thus shares the same orthonormal basis as its associated cell body point.

#### 3 Oral Groove

The *oral groove* is a region on the cell body surface surrounding a curve $\mathbf{G}$ that extends from the anterior toward the cell mouth. We use this curve as a centerline to identify the points belonging to the oral groove. The curve is parameterized by

$$\mathbf{G}(\theta) = \mathbf{X}\big(\theta, \varphi = \gamma(\theta)\big), \tag{8}$$

where $\gamma(\theta)$ specifies the azimuthal location of the groove centerline.
After discretization, we identify body points $\mathbf{x}_{b,i}$ as belonging to the set of oral groove points according to

$$X_{\text{og}} = \{\mathbf{x}_{b,i} \mid \|\mathbf{x}_{b,i} - \mathbf{G}(\theta_i)\| \le w(\theta_i)\}\,, \tag{9}$$

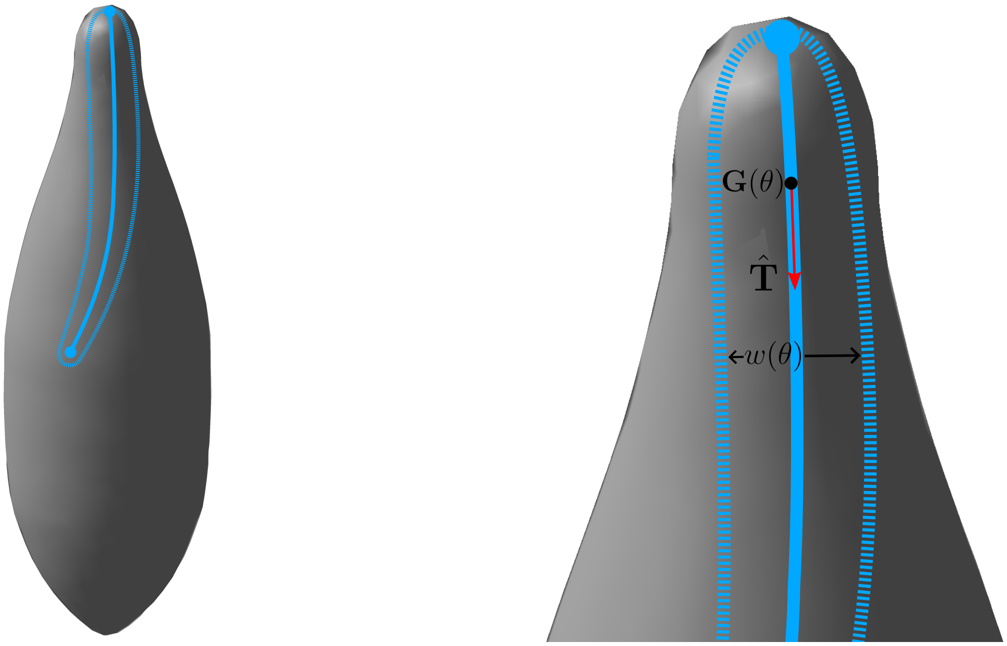


Figure 3: The oral groove on the cell body surface. Mesh points in this region are present but not displayed. (Left) An overview showing the extent of the oral groove. (Right) A close-up of the oral groove with centerline curve, tangent vector, and width function shown.

where $w(\theta)$ specifies the width of the oral groove, which varies along the cell surface. The width function $w(\theta)$ is determined from Figure 4b (main text) [2] by manually selecting points along the boundary of the oral groove and constructing a piecewise cubic Hermite interpolating polynomial through those points.

The cilia within the oral groove beat toward the cell mouth and tangent to the centerline. The tangent vector along the centerline curve is

$$\mathbf{T} = \frac{\partial \mathbf{G}}{\partial \theta}, \tag{10}$$

$$\tag{11}$$

which is normalized to yield

$$\hat{\mathbf{T}} = \frac{\mathbf{T}}{|\mathbf{T}|}. \tag{12}$$

Since $\mathbf{T}$ is a function of $\theta$, each point in $X_{\text{og}}$ has an associated unit tangent vector given by

$$\hat{\mathbf{T}}_i = \frac{\mathbf{T}(\theta_i)}{|\mathbf{T}(\theta_i)|}. \tag{13}$$

### 4 Ciliary Traction Forces

We represent the effect of ciliary activity by prescribing a traction force at each point on the traction layer. This force aims to capture the time-average of the force generated by a collection of cilia in a small region on the cell body. Each force is characterized by a magnitude and a direction,

$$\mathbf{f}_i = A_i \mathbf{V}_i, \tag{14}$$

where $A_i$ is the traction amplitude and $\mathbf{V}_i$ is the corresponding unit direction vector.

We distinguish between two populations of cilia: locomotor cilia covering the cell body, and cilia within the oral groove. These two populations are assigned distinct traction amplitudes and directions tangent to the traction layer.

For the locomotor cilia, the traction forces are directed at an angle $\alpha$ relative to the local $\hat{\theta}$ direction. Thus the prescribed traction forces are

$$\mathbf{f}_{\text{lm},i} = A_{\text{lm}} \left( \cos(\alpha)\hat{\theta}_i + \sin(\alpha)\hat{\varphi}_i \right), \tag{15}$$

where $A_{\text{lm}}$ is the constant traction amplitude of the locomotor cilia.

For the oral groove cilia, the traction forces are directed tangent to the oral groove curve. The prescribed traction is therefore

$$\mathbf{f}_{\text{og},i} = A_{\text{og}} \hat{\mathbf{T}}_i, \tag{16}$$

where $A_{\text{og}}$ is the constant traction amplitude of the oral groove cilia. A diagram of this force distribution is shown in 4.

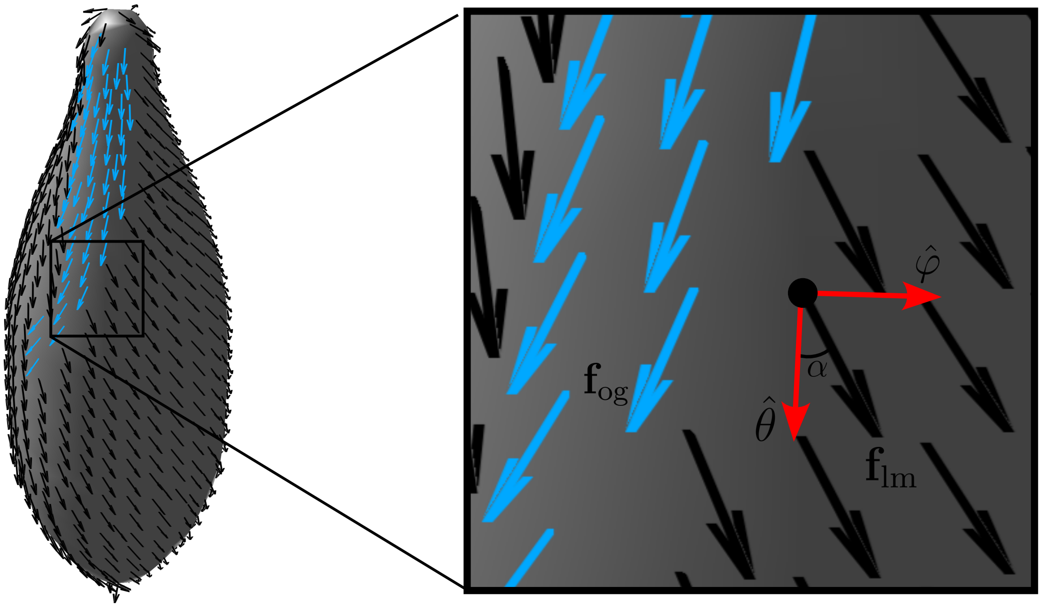


Figure 4: Prescribed forces on the traction layer. The broad view shows the distribution of locomotor ciliary forces (black) and oral groove ciliary forces (blue), with the boxed region indicating the area enlarged in the inset. The close-up illustrates the local force orientations: the oral groove force ($\mathbf{f}_{\text{og}}$) is directed tangent to the groove, i.e. in the $\hat{T}$-direction; the locomotor force ($\mathbf{f}_{\text{lm}}$) is oriented at an angle $\alpha$ relative to the local $\hat{\theta}$ direction. In the simulations presented, $\alpha = 30°$.

#### 5 Cell Orientation in the Lab Frame

The cell body surface, traction layer, and traction forces are defined with respect to the *body frame*, which is centered at the origin and aligned with the cell's long axis along the $z$-axis. We next rotate and translate the surfaces and forces into the *lab frame*, thereby staging our model *Paramecium* above the attachment surface in a prescribed orientation.

To mimic the experimental configuration, the cell is first rotated about its long axis by the azimuthal angle $\phi$. We then apply a combined rotation that brings the cell's long axis from the $z$-axis into the $y$-direction, parallel to the attachment surface ($xy$ plane), and subsequently tilts the cell by an angle of attack

$\beta$. Finally, the cell is translated vertically so that its lowest body points are positioned a ciliary width above the attachment surface.

The rotations are applied in sequence as

$$\mathbf{R} = \mathbf{R}_x \mathbf{R}_z, \tag{17}$$

where

$$\mathbf{R}_z = \begin{pmatrix} \cos(\phi + \pi/2) & -\sin(\phi + \pi/2) & 0 \\ \sin(\phi + \pi/2) & \cos(\phi + \pi/2) & 0 \\ 0 & 0 & 1 \end{pmatrix} \tag{18}$$

rotates the cell about its long axis, and

$$\mathbf{R}_x = \begin{pmatrix} 1 & 0 & 0 \\ 0 & \cos(-\pi/2 - \beta) & -\sin(-\pi/2 - \beta) \\ 0 & \sin(-\pi/2 - \beta) & \cos(-\pi/2 - \beta) \end{pmatrix} \tag{19}$$

both lays the cell onto the attachment surface and applies the prescribed angle of attack. After rotation, the cell is translated in the laboratory $z$-direction such that the lowest point of the cell body surface is a ciliary width above the attachment surface. Thus, body-frame positions and force vectors are transformed according to

$$\mathbf{x}_{\text{lab}} = \mathbf{R}\mathbf{x}_{\text{body}} + \mathbf{x}_0, \qquad \mathbf{f}_{\text{lab}} = \mathbf{R}\mathbf{f}_{\text{body}}, \tag{20}$$

where $\mathbf{x}_0$ is the corresponding vertical translation.

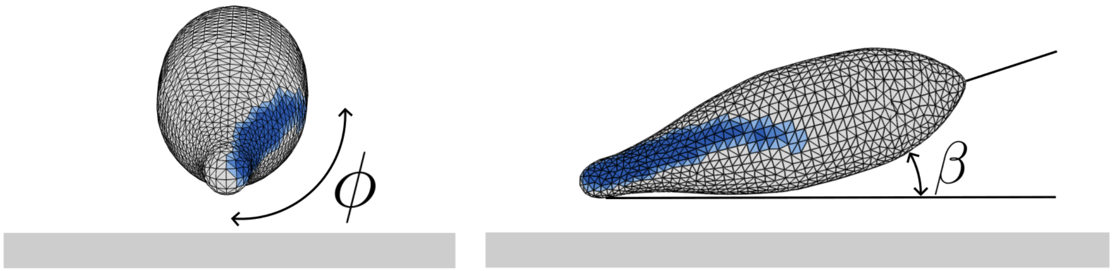


Figure 5: Staging of the cell in the laboratory frame. The cell is rotated from its initial body-frame orientation to reproduce the experimental configuration, with the angle of attack $\beta$ controlling the tilt of the cell relative to the laboratory $xy$ plane and the azimuthal angle $\phi$ controlling its orientation within that plane. The cell is subsequently translated vertically to position the traction layer above the attachment surface. Here, $\beta = 15°$ and $\phi = 90°$. In the simulations presented, $\|\mathbf{x}_0\| = 10\,\mu\text{m}$.

### 6 Flow Computation

The forces on the cell body and the surrounding fluid velocity are computed using the method of images for regularized Stokeslets (MIRS). This method provides a regularized fundamental solution to the Stokes equations that accounts for the presence of a plane wall where the velocity is zero. The details of the MIRS formulation are described in [1].

For a traction force $\mathbf{f}_i$ applied at a source point $\mathbf{x}_i$, the velocity induced at a target point $\mathbf{x}$ is given by

$$\mathbf{u}(\mathbf{x}) = \mathbf{S}^{\epsilon}(\mathbf{x}, \mathbf{x}_i)\mathbf{f}_i \Delta A_i, \tag{21}$$

where $\mathbf{S}^{\epsilon}(\mathbf{x}, \mathbf{x}_i)$ is the tensor called the *regularized Stokeslet*, $\epsilon$ is the regularization parameter, and $\Delta A_i$ is the area of the surface element assigned to the point $\mathbf{x}_i$. In our model we have collections of source points ($X_b$, $X_t$) and point forces ($\mathbf{f}_b$, $\mathbf{f}_t$). By the linearity of the Stokes equations, the velocity at a target point arising from these collections is given by

$$\mathbf{u}(\mathbf{x}) = \sum_{i=1}^{N} \mathbf{S}^{\epsilon}(\mathbf{x}, \mathbf{x}_{b,i})\mathbf{f}_{b,i} \Delta A_i^b + \sum_{i=1}^{N} \mathbf{S}^{\epsilon}(\mathbf{x}, \mathbf{x}_{t,i})\mathbf{f}_{t,i} \Delta A_i^t. \tag{22}$$

For a collection of target points, such as the points on a spatial grid, this relation can be written in matrix form as

$$\mathcal{U} = \mathcal{M}_b \mathcal{F}_b + \mathcal{M}_t \mathcal{F}_t. \tag{23}$$

where $\mathcal{U}$ is the collection of velocities at the target points, $\mathcal{F}_b$ and $\mathcal{F}_t$ are the collections of forces on the cell body and ciliary traction layer, respectively, and $\mathcal{M}_b$ and $\mathcal{M}_t$ are the corresponding MIRS mobility matrices.

The ciliary forces $\mathcal{F}_t$ are prescribed from the traction model described above. The cell body forces $\mathcal{F}_b$, however, are unknown. Because the cell is stationary in the lab frame, the velocity at every cell body point must satisfy

$$\mathcal{U}_b = \mathbf{0}. \tag{24}$$

Evaluating the matrix relation at the cell body points gives

$$\mathbf{0} = \mathcal{M}_b \mathcal{F}_b + \mathcal{M}_t \mathcal{F}_t, \tag{25}$$

which is solved for the unknown body forces $\mathcal{F}_b$. Once $\mathcal{F}_b$ has been determined, the complete set of forces is known and the velocity at any collection of target points can be computed from Equation 23.

### 7 Streamline Visualization

For visualization, we evaluate the resulting velocity field on Cartesian grids and use the computed velocities to generate streamlines. We consider two primary visualizations of the flow.

For the two-dimensional flow visualization in Figure 6 (main text), we first construct a uniform two-dimensional Cartesian grid in a plane at a fixed height above the attachment surface, then solve for the three-dimensional velocity field at each grid point. We then take the in-plane components of the velocity, $u_x$ and $u_y$, to obtain a two-dimensional velocity field. Streamlines of this field are subsequently generated and plotted.

For the three-dimensional flow visualizations in Figures 6 and 7, we instead construct a three-dimensional Cartesian grid surrounding the cell and evaluate all three components of the velocity field throughout the domain. Streamline seeds are distributed on a plane between the cell and the attachment surface. From each seed point, streamlines are integrated in both the forward and reverse directions through the three-dimensional velocity field, providing a visualization of the local flow structure around the staged cell.

### 8 Varying the Cell Orientation

By varying the cell orientation, we can discern which flow features are attributable to cell orientation vs. ciliary force distributions. In Figure 7, we show computed flow fields for four orientations of the cell. The typical posture observed in Figures 4 and S1 (main text) is showcased in 7b.

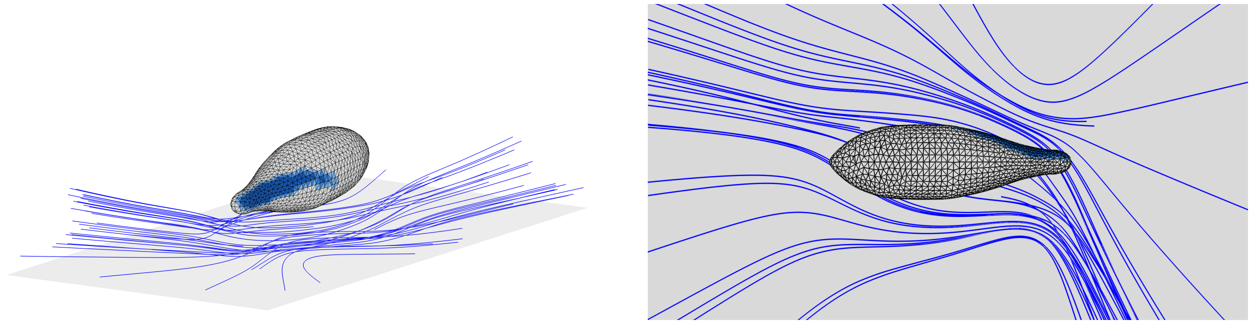

Figure 6: Visualizations of the three-dimensional flow field around the staged *Paramecium*. (Left) A perspective view. Streamlines are seeded in a plane between the cell and the attachment surface and integrated in both the forward and reverse directions through the computed three-dimensional velocity field. (Right) A projected view. The flow field is viewed from above the attachment surface, as in Figure 6 (main text). Here, $\phi = 90°$, $\beta = 5°$, and $A_{\mathrm{lm}}/A_{\mathrm{og}} = 0.05$.

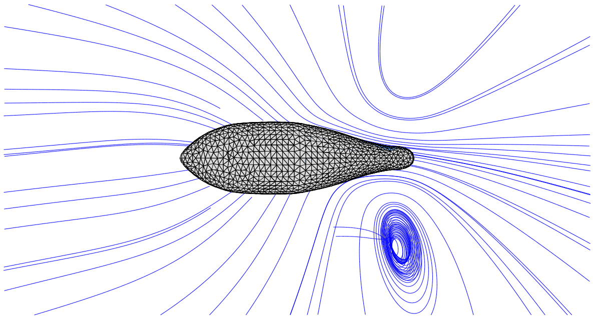

(a) $\phi = 0°$

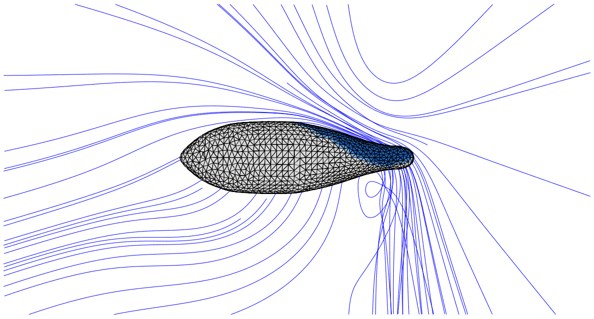

(b) $\phi = 90°$

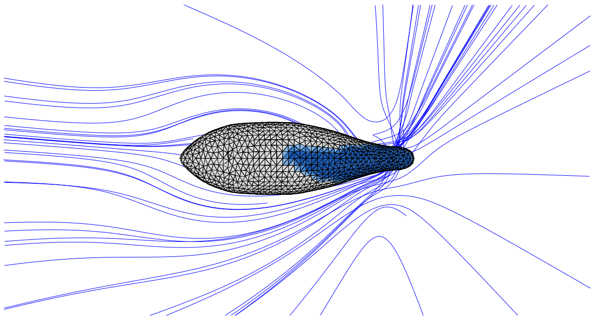

(c) $\phi = 180°$

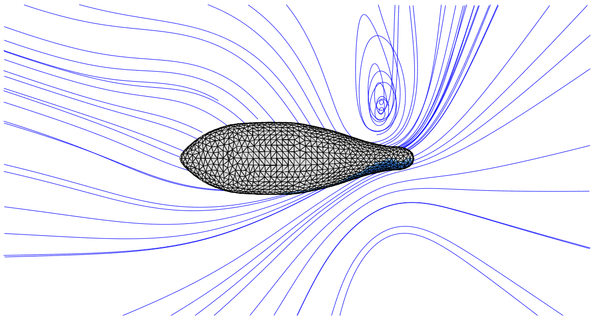

(d) $\phi = 270°$

Figure 7: Computed flow fields for four orientations of the cell. Streamlines are computed from the full three-dimensional flow, and viewed from above the attachment surface in projection. Here, $\phi = 90°$, $\beta = 5°$, and $A_{\mathrm{lm}}/A_{\mathrm{og}} = 0.05$.

### References

[1] Josephine Ainley, Sandra Durkin, Rafael Embid, Priya Boindala, and Ricardo Cortez. The method of images for regularized Stokeslets. *Journal of Computational Physics*, 227(9):4600–4616, April 2008.

[2] S. O. Mast. The Food-Vacuole in *Paramecium*. *The Biological Bulletin*, 92(1):31–72, February 1947.

[3] Per-Olof Persson and Gilbert Strang. A Simple Mesh Generator in MATLAB. *SIAM Review*, 46(2):329–345, January 2004.
